# Determinants of domain-specific cognitive impairment after stroke: a mega-analysis of 13 acute ischaemic stroke cohorts

**DOI:** 10.64898/2026.11.10.765579

**Authors:** Marvin Petersen, Nick A. Weaver, Kaustubh R. Patil, Hugo P. Aben, Hee-Joon Bae, Jonathan G. Best, J. Matthijs Biesbroek, Régis Bordet, Francesca Chappell, Mirthe Coenen, Thibaut Dondaine, Jule Filler, Michael D. Fox, Olivier Godefroy, Saima Hilal, Andreas Horn, Yeonwook Kang, Jonguk Kim, Sebastian Koehler, Floor A.S. de Kort, Paul L.M. de Kort, Mi Sun Oh, Bonnie Y.K. Lam, Byung-Chul Lee, Keon-Joo Lee, Minwoo Lee, Jae-Sung Lim, Renaud Lopes, Stephen D.J. Makin, Vincent C.T. Mok, Martine Roussel, Frederic L.W.V.J. Schaper, Julie Staals, An Qi Toh, Maria C. Valdés Hernández, Ruben S. van der Giessen, Narayanaswamy Venketasubramanian, David J. Werring, Joanna M. Wardlaw, Adrian Wong, Xin Xu, Kyung-Ho Yu, Christopher P.L.H. Chen, Simon B. Eickhoff, Geert Jan Biessels, Meta VCI Map consortium

## Abstract

The relationship between brain lesions and cognition is enigmatic. Apparently similar lesions can produce different outcomes. Stroke provides a prototypical model to understand how lesion anatomy, network disruption, prior injury and patient characteristics shape impairment. We analysed 3,034 patients from 13 acute ischaemic stroke cohorts, integrating infarct size and location, structural and functional lesion network mapping, pre-existing cerebrovascular injury, and demographic and clinical characteristics to predict impairment across six cognitive domains within one year after stroke. Infarct topography and connectivity improved prediction for six of seven outcomes over demographic, clinical and infarct-size information alone. Structural lesion connectivity accounted for most gains, with additional gains from topography and connectivity of pre-existing vascular lesions. Across domains, gains rested on left anterior thalamic connectivity plus domain-specific anatomy. Our findings establish, at unprecedented scale, that understanding lesion–cognition relationships requires looking beyond the lesion to networks, prior brain pathology and individual context.

## Background

Focal lesions are neuroscience’s oldest source of evidence that a brain region is necessary for a cognitive function, not merely involved in it.^1,2^ Yet the relationship between brain lesions and cognitive deficits is enigmatic. Patients with lesions in similar locations, sometimes even of comparable size, often exhibit markedly different cognitive outcomes. This variability likely reflects factors beyond local brain tissue loss: disruption of the distributed brain systems the lesion affects and the biological context in which it occurs, such as brain reserve and pre-existing brain injury.^3,4^ Explaining cognitive outcome after focal brain injury therefore requires understanding these factors together.

Ischaemic stroke provides a prototypical model for studying lesion-deficit relationships. Brain infarcts are usually focal, have a defined onset, and are routinely characterised with neuroimaging, enabling large-scale investigations. Post-stroke cognitive impairment (PSCI) is a common consequence of stroke, affecting more than half of survivors, substantially reducing quality of life and functional independence.^5^ Infarct characteristics like location and size are important determinants of cognitive impact.^6^ However, current location and size-based models insufficiently explain occurrence of PSCI. Lesion network mapping (LNM) captures the broader network context of focal brain injury, showing that disruption of distributed brain circuitry explains domain-specific cognitive deficits beyond the local consequences of the lesion itself.^7–10^ Moreover, pre-existing cerebrovascular injury, commonly present in patients with stroke, has emerged as another independent determinant of PSCI.^11^ Finally, patient-level characteristics, including demographics, vascular risk factors and cognitive ability earlier in life, further contribute to cognitive outcome.^12^ Collectively, these observations suggest that post-stroke cognition is shaped by the interplay of three dimensions: the infarct and the networks it disrupts, the prior disease burden of the brain in which it occurs, and the characteristics of the individual who sustains it. Understanding their relative and joint contributions to cognitive outcome therefore requires an analytical framework that integrates these dimensions.

Predictive modelling provides such a framework.^13^ Models are trained on one set of patients and evaluated on unseen patients, as a prognostic model would be used in clinical practice. Repeating this procedure across multiple data partitions assesses the robustness and generalisability of model performance. Because models can be trained on any chosen set of features, each dimension – infarct size, location, and network connectivity, pre-existing injury, demographics and clinical characteristics – can be modelled on its own and then integrated, to ask how much each adds to predicting cognitive outcomes in unseen patients beyond the others. Importantly, improved prediction accuracy alone is not equivalent to biological understanding. Predictive performance assessments must therefore be accompanied by analyses that link any model improvement to interpretable anatomical profiles corresponding to biologically meaningful brain organisation, and demonstrate that it generalises across independent patient populations, a prerequisite for any prognostic use.^13–15^

Here, we implement a mega-analysis of 3,034 patients from 13 cohorts of the Meta VCI Map consortium with harmonised domain-specific cognitive outcomes assessed up to one year after stroke, integrating complementary determinants of PSCI within a unified predictive framework. This framework trains models to jointly evaluate demographic and clinical characteristics, acute infarct location, functional and structural LNM, and pre-existing cerebrovascular injury. Specifically, we test whether combining acute infarct anatomy, pre-existing cerebrovascular injury, demographics and clinical information improves prediction of domain-specific PSCI beyond a clinical baseline model, and assess the contribution of each. We then examine whether the improvement holds across varying sample sizes, modelling choices, and independent patient cohorts. Next, we examine the anatomy underlying prediction: whether the circuits the models rely on are shared across cognitive domains or specific to each, and whether patients, grouped by their infarct connectivity, form anatomically coherent subgroups with distinct cognitive risk profiles. Finally, we extend the framework to pre-existing cerebrovascular injury, to determine whether white matter hyperintensities (WMH), old infarcts and lacunes carry outcome information beyond the acute infarct, also considering their own topography and connectivity.

## Results

Patients were on average 66.8 years old (SD ± 11.6), 39.0% were female, and the median NIHSS score at admission was 3 (IQR 1–5). The median acute infarct volume was 2.9 mL (IQR 1.0–14.7); 58.5% of infarcts were smaller than 5 mL, 27.9% between 5 mL and 30 mL, and 13.6% were 30 mL or greater. Cognitive assessment was performed a median of 105 days after the index stroke (IQR 74–170). Outcomes were defined as binary impairment status: a cognitive domain was classified as impaired for a patient when more than half of its tests fell below the fifth percentile of local normative data, and any-domain PSCI as impairment in at least one domain. Any-domain PSCI was present in 43.0% of patients, and domain-specific impairment ranged from 11.0% for visuospatial memory to 26.7% for visuospatial function. Sample and cohort characteristics are given in *table 1* and *supplementary table S1*. The spatial distribution of acute infarcts is shown in *supplementary figure S1*. Unless stated otherwise, lesion refers to the acute infarct; pre-existing cerebrovascular lesions (WMH, old infarcts and lacunes) are named as such.

**Table 1.**
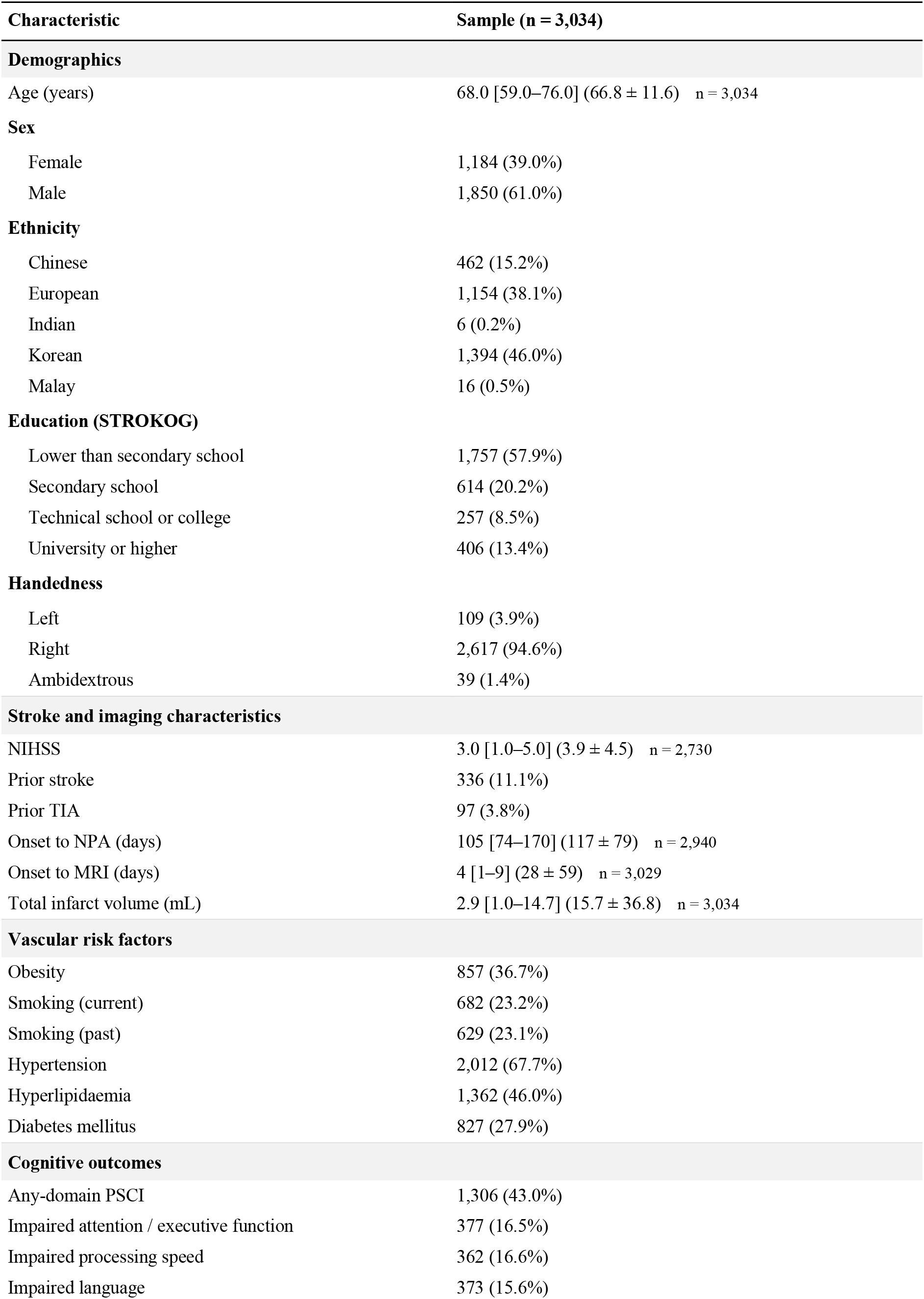
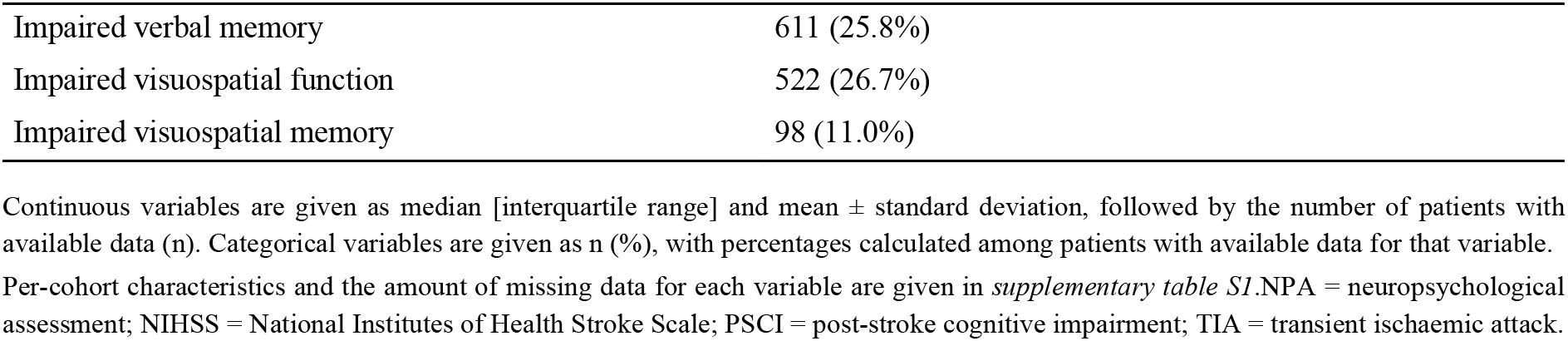
Characteristics of the study population.

| Characteristic | Sample (n = 3,034) |
|---|---|
| **Demographics** |  |
| Age (years) | 68.0 [59.0–76.0] (66.8 ± 11.6) n = 3,034 |
| **Sex** |  |
| Female | 1,184 (39.0%) |
| Male | 1,850 (61.0%) |
| **Ethnicity** |  |
| Chinese | 462 (15.2%) |
| European | 1,154 (38.1%) |
| Indian | 6 (0.2%) |
| Korean | 1,394 (46.0%) |
| Malay | 16 (0.5%) |
| **Education (STROKOG)** |  |
| Lower than secondary school | 1,757 (57.9%) |
| Secondary school | 614 (20.2%) |
| Technical school or college | 257 (8.5%) |
| University or higher | 406 (13.4%) |
| **Handedness** |  |
| Left | 109 (3.9%) |
| Right | 2,617 (94.6%) |
| Ambidextrous | 39 (1.4%) |
| **Stroke and imaging characteristics** |  |
| NIHSS | 3.0 [1.0–5.0] (3.9 ± 4.5) n = 2,730 |
| Prior stroke | 336 (11.1%) |
| Prior TIA | 97 (3.8%) |
| Onset to NPA (days) | 105 [74–170] (117 ± 79) n = 2,940 |
| Onset to MRI (days) | 4 [1–9] (28 ± 59) n = 3,029 |
| Total infarct volume (mL) | 2.9 [1.0–14.7] (15.7 ± 36.8) n = 3,034 |
| **Vascular risk factors** |  |
| Obesity | 857 (36.7%) |
| Smoking (current) | 682 (23.2%) |
| Smoking (past) | 629 (23.1%) |
| Hypertension | 2,012 (67.7%) |
| Hyperlipidaemia | 1,362 (46.0%) |
| Diabetes mellitus | 827 (27.9%) |
| **Cognitive outcomes** |  |
| Any-domain PSCI | 1,306 (43.0%) |
| Impaired attention / executive function | 377 (16.5%) |
| Impaired processing speed | 362 (16.6%) |
| Impaired language | 373 (15.6%) |
| Impaired verbal memory | 611 (25.8%) |
| Impaired visuospatial function | 522 (26.7%) |
| Impaired visuospatial memory | 98 (11.0%) |

Continuous variables are given as median [interquartile range] and mean ± standard deviation, followed by the number of patients with available data (n). Categorical variables are given as n (%), with percentages calculated among patients with available data for that variable.
Per-cohort characteristics and the amount of missing data for each variable are given in *supplementary table S1*.NPA = neuropsychological assessment; NIHSS = National Institutes of Health Stroke Scale; PSCI = post-stroke cognitive impairment; TIA = transient ischaemic attack.

A study overview is provided in *figure 1*.

**Figure 1.**
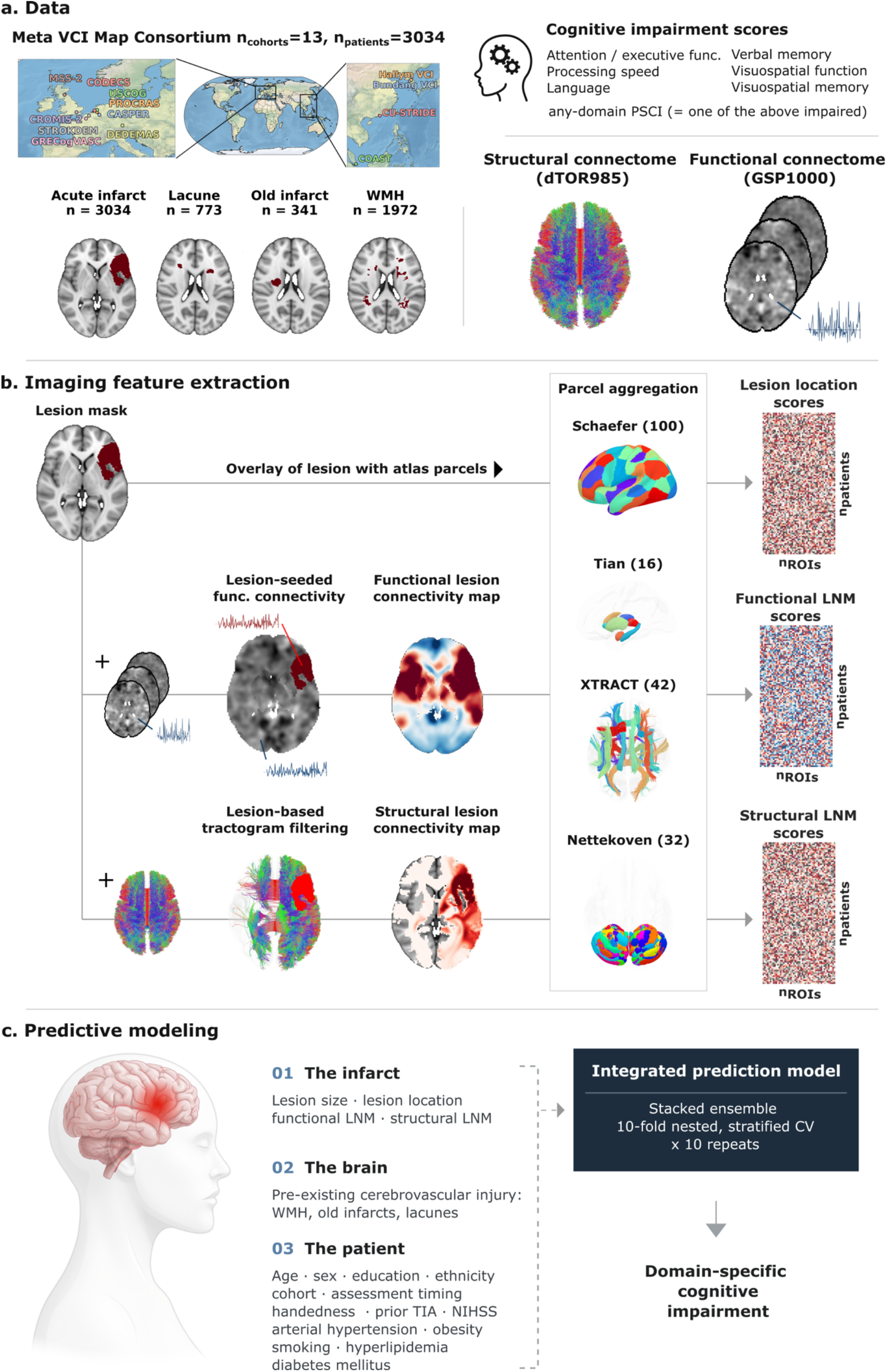
Study overview. **(a)** Data. Individual-patient data were pooled from 13 cohorts of the Meta VCI Map consortium (n = 3,034 patients with acute ischaemic stroke). Pre-existing lesions were segmented in a subset of 1,973 patients, in whom WMH were present in 1,972, lacunes in 773 and old infarcts in 341; acute infarcts were available for all 3,034 patients. Cognitive outcomes comprised binary impairment labels for six domains (attention/executive function, processing speed, language, verbal memory, visuospatial function, visuospatial memory) and a composite any-domain PSCI indicator. Two normative connectomes served as the basis for network mapping: a population-level structural tractogram (dTOR985) and a resting-state functional connectome (GSP1000). **(b)** Imaging feature extraction. For each patient, the binary lesion mask was processed through three parallel pathways. First, the lesion mask was directly overlaid on the atlas parcellation to compute per-parcel lesion location scores (percent parcel damage). Second, the lesion mask was combined with the GSP1000 functional connectome to produce a voxelwise functional connectivity map, which was then aggregated to per-parcel mean *t*-statistics (functional LNM scores). Third, the lesion mask was used to filter the dTOR985 tractogram, and the proportion of lesion-interfacing streamlines per parcel was computed (structural LNM scores). All three feature vectors were aggregated over the same atlas regimen comprising four complementary parcellations: Schaefer (cortex, 100 parcels), Tian (subcortex, 16 parcels), XTRACT (white matter, 42 tracts), and Nettekoven (cerebellum, 32 parcels), yielding 190 features per modality per patient. **(c)** Feature sets were organised into three groups: the infarct, represented by lesion size, lesion location, functional LNM and structural LNM scores; the brain in which it occurred, represented by pre-existing cerebrovascular injury, including WMH, old infarcts and lacunes, and characterised by lesion location, functional LNM and structural LNM scores; and the patient, represented by age, sex, education, ethnicity, cohort, interval from stroke onset to cognitive assessment, interval from stroke onset to MRI, handedness, prior TIA, NIHSS, and vascular risk factors, including arterial hypertension, obesity, smoking, hyperlipidaemia and diabetes mellitus. These were combined in an integrated model to predict impairment in each cognitive domain and any-domain PSCI. CV = cross-validation; dTOR985 = Toronto diffusion-weighted structural connectome (N = 985); GSP1000 = Brain Genomics Superstruct Project functional connectome (N = 1000); functional LNM = functional lesion network mapping; MRI = magnetic resonance imaging; NIHSS = National Institutes of Health Stroke Scale; PSCI = post-stroke cognitive impairment; structural LNM = structural lesion network mapping; TIA = transient ischaemic attack; WMH = white matter hyperintensities of presumed vascular origin.

### Integrating acute infarct anatomy, demographics and clinical characteristics improved prediction of post-stroke cognitive impairment

Models were evaluated by 10 × 10 repeated stratified cross-validation of the pooled sample, with performance quantified as the area under the precision–recall curve (AUPRC), a metric suited to imbalanced outcomes i.e. binary outcomes in which one class is rarer, as impairment was in most domains. We compared four models – two null, one baseline and the integrated model. The two null models defined chance-level performance: a majority-class model, whose AUPRC equals impairment prevalence, and a model trained on permuted outcome labels, which verifies that the modelling pipeline does not extract signal from noise. The baseline model comprised demographic and clinical characteristics and infarct volume. The integrated model added three sets of acute infarct scores, each computed for 190 predefined brain regions (for details see *figure 1b* and *Methods*): lesion location (the proportion of each region damaged), functional LNM (each region’s functional connectivity to the infarct) and structural LNM (the proportion of each region’s streamlines passing through the infarct). Each feature set was modelled by a separate ridge base learner, and their predictions were combined through linear stacking.

To attribute gains to individual feature sets, we additionally evaluated models adding each acute infarct feature set to the baseline alone, and models removing each from the integrated model. Subsequent model comparisons are reported as the difference in AUPRC (ΔAUPRC) between a model and its stated reference, computed on identical folds.

The baseline model predicted all seven cognitive targets significantly better than both null models. The integrated model further improved prediction significantly beyond the baseline for six of seven targets (FDR-corrected Nadeau–Bengio *P* < 0.05; *figure 2a*). For any-domain PSCI, AUPRC increased from 0.43 and 0.44 for the majority-class and permuted models to 0.67 for the baseline and 0.71 for the integrated model. Gains in the cognitive domains (ΔAUPRC, integrated minus baseline) ranged from 0.02 to 0.08 (baseline → integrated: attention/executive function 0.43 → 0.48, processing speed 0.43 → 0.45, language 0.42 → 0.47, verbal memory 0.49 → 0.57, visuospatial function 0.47 → 0.51), with verbal memory benefiting most. Visuospatial memory, assessed in the smallest subsample (n = 889), was least well predicted, and the integrated model did not significantly improve on the baseline, although it remained above both null models (0.27 → 0.28). AUROC results were consistent (*supplementary figure S2*). Predicted probabilities of the integrated model were correct on average for every target (calibration-in-the-large between −0.005 and 0.004). Calibration slopes were 1.04 for any-domain PSCI and above 1 for all cognitive domains, indicating that predictions were conservative rather than overconfident (*supplementary figure S3*).

**Figure 2.**
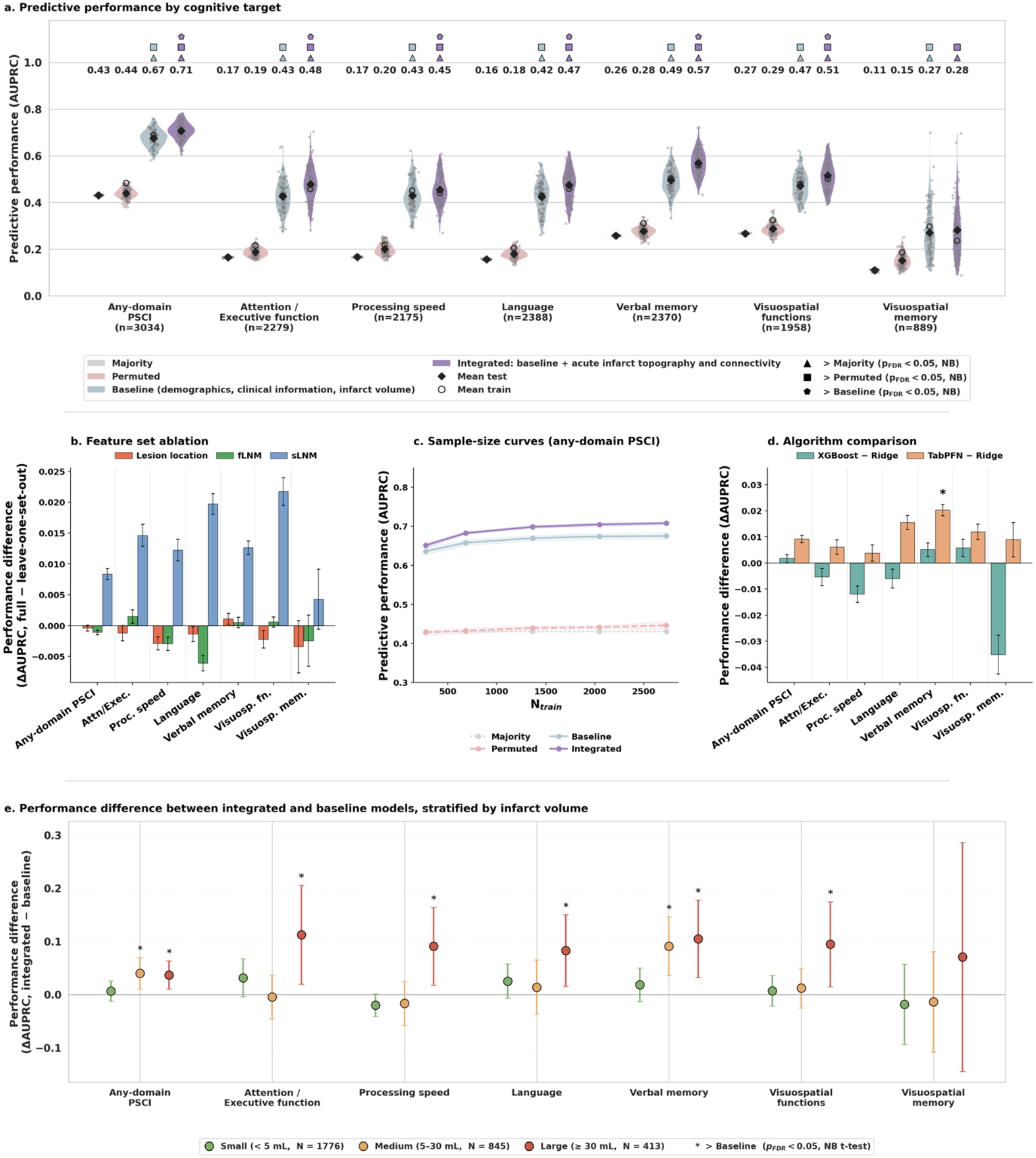
Prediction of PSCI based on acute infarct topography and connectivity, demographics and clinical information. **(a)** Predictive performance for each cognitive target based on 10 × 10 repeated stratified cross-validation. Four models were assessed: majority (model always predicting prevalence), permuted (model trained on shuffled labels), baseline (demographics, clinical information, and infarct volume), and integrated (baseline plus multi-modal acute infarct topography: lesion location, functional LNM, and structural LNM). Violins and grey dots show the distribution of test AUPRC across the 100 outer folds (10 repeats × 10 folds); diamonds indicate mean test AUPRC across folds, with the value printed above each model’s violin; open circles indicate mean train AUPRC across folds. Symbols above the violins indicate that the model performed significantly better than a reference model (FDR-corrected Nadeau–Bengio paired t-tests, *p*_FDR_ < 0.05): triangle, better than majority; square, better than permuted; pentagon, better than baseline. The integrated model performed significantly better than the baseline on six of the seven cognitive targets. **(b)** To quantify the unique contribution of each lesion anatomy representation, each lesion feature set was removed in turn from the integrated model, which was then re-fitted on the remaining features using the same outer folds. ΔAUPRC (full model minus drop-one-feature-set model) is shown per cognitive target. Error bars denote across-fold SE. Ablating structural LNM produced the largest performance drop for all seven cognitive targets, whereas removing lesion location or functional LNM had negligible effects. **(c)** Sample-size curve for any-domain PSCI. Mean predictive performance (AUPRC ± SE across folds) is plotted as a function of training set size (N_train_) for each model. Both the baseline and integrated models approached a plateau at larger training set sizes, with the integrated model converging at a higher AUPRC, while the majority and permuted references remained flat at their null floors. **(d)** Algorithm comparison. Difference in mean test AUPRC (ΔAUPRC ± SE across folds) of the stacked ensemble with XGBoost or TabPFN imaging base learners relative to the ridge-based ensemble, under the same CV architecture and shown for each cognitive target. Differences were small; TabPFN reached the highest point estimates for all cognitive targets. Asterisks mark significant improvement over ridge (FDR-corrected Nadeau–Bengio paired t-test, *p*_FDR_ < 0.05). **(e)** Predictive performance difference (ΔAUPRC) of the integrated over baseline model, stratified by infarct volume (small: <5 mL; medium: 5–30 mL; large: ≥30 mL). Points show the mean fold-paired difference across the 100 outer folds within each stratum; whiskers denote the 95% confidence intervals based on Nadeau-Bengio corrected variance. Asterisks denote integrated > baseline (one-sided NB *t*-tests, FDR-corrected across the three volume strata within each cognitive target, *p*_FDR_ < 0.05). The predictive performance gain from infarct topography and connectivity over the baseline was largest and most consistently significant in patients with large infarcts, moderate for medium infarcts, and low for small infarcts. AUPRC = area under the precision–recall curve; CV = cross-validation; functional LNM = functional lesion network mapping; structural LNM = structural lesion network mapping; NB = Nadeau–Bengio; FDR = false discovery rate; SE = standard error.

### Structural lesion connectivity of the acute infarct accounted for most of the improvement beyond baseline

When each acute infarct feature set was added to the baseline model individually, structural LNM achieved the highest performance for all seven cognitive targets, while lesion location and functional LNM each improved on the baseline for three of seven (*supplementary figure S4*). Conversely, removing the structural LNM scores from the integrated model caused the largest drops for all seven cognitive outcomes (ΔAUPRC 0.004 to 0.022), whereas removing lesion location or functional LNM scores had little effect (*figure 2b*). Within the baseline model, ablation of individual variables showed that cohort membership, NIHSS, infarct volume, and time between stroke onset and neuropsychological assessment contributed most strongly to predictive performance (*supplementary figure S5*).

### The predictive benefit was stable across sample size and modelling algorithm

Learning curves were obtained by training on random subsamples of increasing size drawn from each training fold. The advantage of the integrated model emerged early, at roughly 20% of the training data, and widened with sample size before levelling off towards the full dataset (*figure 2c*).

To test whether linear modelling of the input features was limiting performance, we replaced the ridge base learners with two more flexible algorithms, TabPFN, a tabular foundation model, and XGBoost, a nonlinear tree-based ensemble, while keeping the stacking architecture unchanged. XGBoost did not perform significantly better than the linear model for any cognitive outcome, and TabPFN did so only for verbal memory (ΔAUPRC = +0.020, Nadeau–Bengio *p*_FDR_ = 0.030; *figure 2d*), although it reached the highest point estimates for every cognitive outcome. Given these small differences, we retained the linear stacking model as an interpretable default.

### The benefit scaled with infarct size and was present across age and sex

Because most infarcts in our sample were small, we examined whether the benefit of detailed lesion characterisation over the baseline depended on infarct size (*figure 2e*). Averaged across the seven cognitive targets, the benefit was small for infarcts <5 mL (mean ΔAUPRC ± SD = +0.008 ± 0.020; median volume = 1.2 mL), modest for medium-sized infarcts of 5–30 mL (+0.018 ± 0.038; 12.8 mL), and greatest for infarcts ≥30 mL (+0.085 ± 0.025; 57.5 mL).

In analyses stratified by age, the integrated model improved on the baseline in every age tertile, but the pattern across cognitive targets was heterogeneous. Mean gains across targets were similar in the youngest (<63 years; ΔAUPRC +0.026) and middle tertiles (63 to <73 years; +0.026) and smaller in the oldest tertile (≥73 years; +0.008) (*supplementary figure S6*). In analyses stratified by sex, the point estimate of the gain was larger in women than in men for six of seven targets, all except visuospatial memory, and gains were significant for five targets in women and three in men (*supplementary figure S7*).

### Alternative lesion feature derivations did not improve prediction

Finer parcellations slightly reduced the performance of the integrated model for six of seven cognitive targets (*supplementary figure S8*). Subsampling the normative tractogram to as few as 10% of its streamlines left performance unchanged (*supplementary figure S9*). Entering log-transformed instead of raw infarct volume left the performance of the integrated model and its significant advantage over baseline unchanged (supplementary figure S10). Replacing regional scores by principal components of the voxelwise maps resulted in equal or lower mean AUPRC and AUROC for every lesion representation and cognitive target (*supplementary figure S11*). The 190-region reference parcellation, regional feature aggregation, the full normative tractogram and raw infarct volume are therefore adequate defaults for this application.

### Predictive gains extended to independent stroke cohorts

Because pooled cross-validation tests patients from the cohorts seen during training, we repeated the analysis under leave-one-cohort-out cross-validation to test transfer to unseen cohorts (*supplementary figures S12 to S14*). Although predictive performance decreased relative to pooled cross-validation, the integrated model continued to perform better than the baseline model for three of seven cognitive targets based on normalised AUPRC (any-domain PSCI, language and verbal memory) and for four of seven based on AUROC (additionally processing speed; FDR-corrected Wilcoxon P < 0.05; *supplementary figure S12a and S13*). When each cohort was held out in turn, the integrated model predicted any-domain PSCI better than the baseline in all 13 (*supplementary figure S12b*). Absolute performance differed widely between cohorts and was lowest in CODECS and DEDEMAS. Cross-cohort performance of the integrated model tended to be higher in larger cohorts and in cohorts with more severe strokes and lower in cohorts with higher median education or more missing covariate data (*supplementary figure S12c and S12d*). Performance of individual lesion representations under leave-one-cohort-out cross-validation is shown in *supplementary figure S15*.

### Structural lesion network mapping reveals biologically meaningful neuroanatomical profiles

To unravel the anatomy underlying the improved prediction, we examined the structural LNM base learner, as structural LNM was the most informative of the three acute infarct feature sets. Averaging its coefficients across cross-validation folds and retaining regions with stable coefficients identified the regions the model relied on to predict impairment. Structural connectivity of the infarct to the left anterior thalamus was the most consistent predictor: it carried a stable positive coefficient for all seven targets and ranked among the five most strongly weighted regions for five of them (any-domain PSCI, processing speed, language, verbal memory and visuospatial function). The adjacent left anterior thalamic radiation ranked among the top five for any-domain PSCI, language and verbal memory. Superimposed on this shared core were domain-specific profiles (*figure 3*): 17 of the 24 regions that ranked among the top five for any target did so for a single target only (*supplementary figure S16*). Coefficient profiles of the lesion location and functional LNM base learners are shown in *supplementary figures S17* and *S18*.

**Figure 3.**
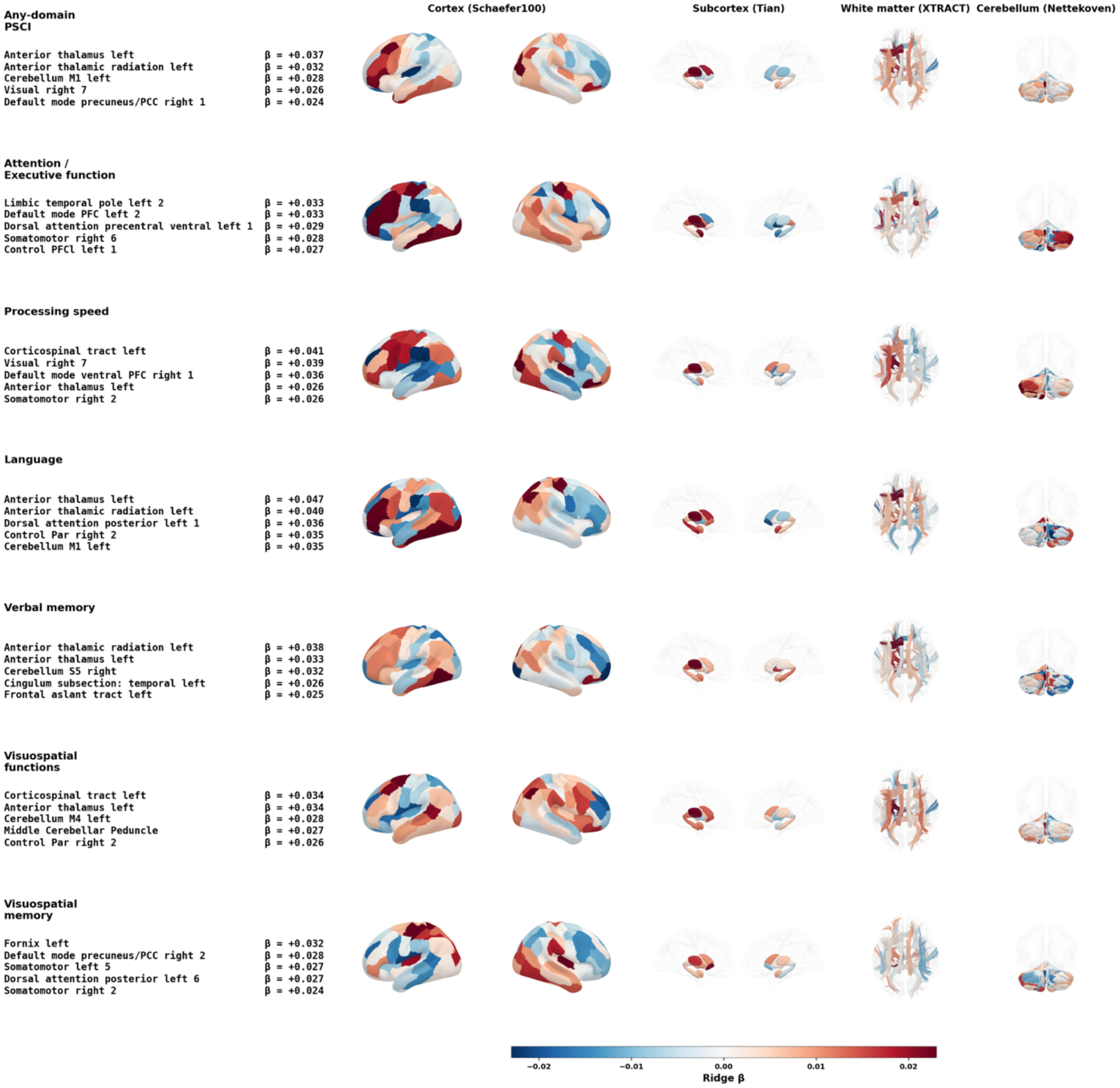
Anatomy of cognitive domain-specific structural lesion network profiles. For each cognitive target, standardised β coefficients were extracted from the base learner informed by structural LNM. The coefficients were averaged across repeated cross-validation folds and displayed across cortical (Schaefer), subcortical (Tian), white matter (XTRACT), and cerebellar (Nettekoven) brain regions. Positive values (red) indicate that the model weights greater structural connectivity of a region to the lesion towards a higher predicted probability of impairment; negative values (blue) indicate the opposite weighting. For each cognitive domain, the five regions with the largest positive β values are listed at the far left. Structural lesion connectivity of the left anterior thalamus and left anterior thalamic radiation were among the most consistent predictors across cognitive domains. At the same time, each domain showed additional, more specific profiles of structural lesion connectivity. SE = standard error.

We next examined whether structural lesion connectivity alone, without reference to cognitive outcome, organised patients into anatomically coherent subgroups, and how cognitive risk was distributed across them. We embedded patient-level structural LNM scores using UMAP and clustered them with HDBSCAN. This identified eight clusters corresponding to canonical vascular territories – left and right MCA, left and right deep perforator, left and right PCA/AChA, ACA and infratentorial – and a ninth, anatomically heterogeneous cluster (*figure 4a and d*). Any-domain PSCI prevalence ranged from 31% to 67% across clusters and was highest for left PCA/AChA (67%) and ACA (64%) infarcts, exceeding the MCA territories (52% and 47%) despite their larger infarcts. In addition, any-domain PSCI prevalence was higher for the left than the right cluster of every paired territory. Domain-specific predictions followed lateralised patterns: left MCA and left deep perforator clusters had higher predicted probabilities of impairment in attention/executive function, language and verbal memory, and right-hemispheric clusters in visuospatial function (*figure 4b, c, e and f*). For the distribution of baseline variables on the UMAP, see *supplementary figure S19*. Embedding and clustering results for lesion location and functional LNM are presented in *supplementary figures S20* and *S21*.

**Figure 4.**
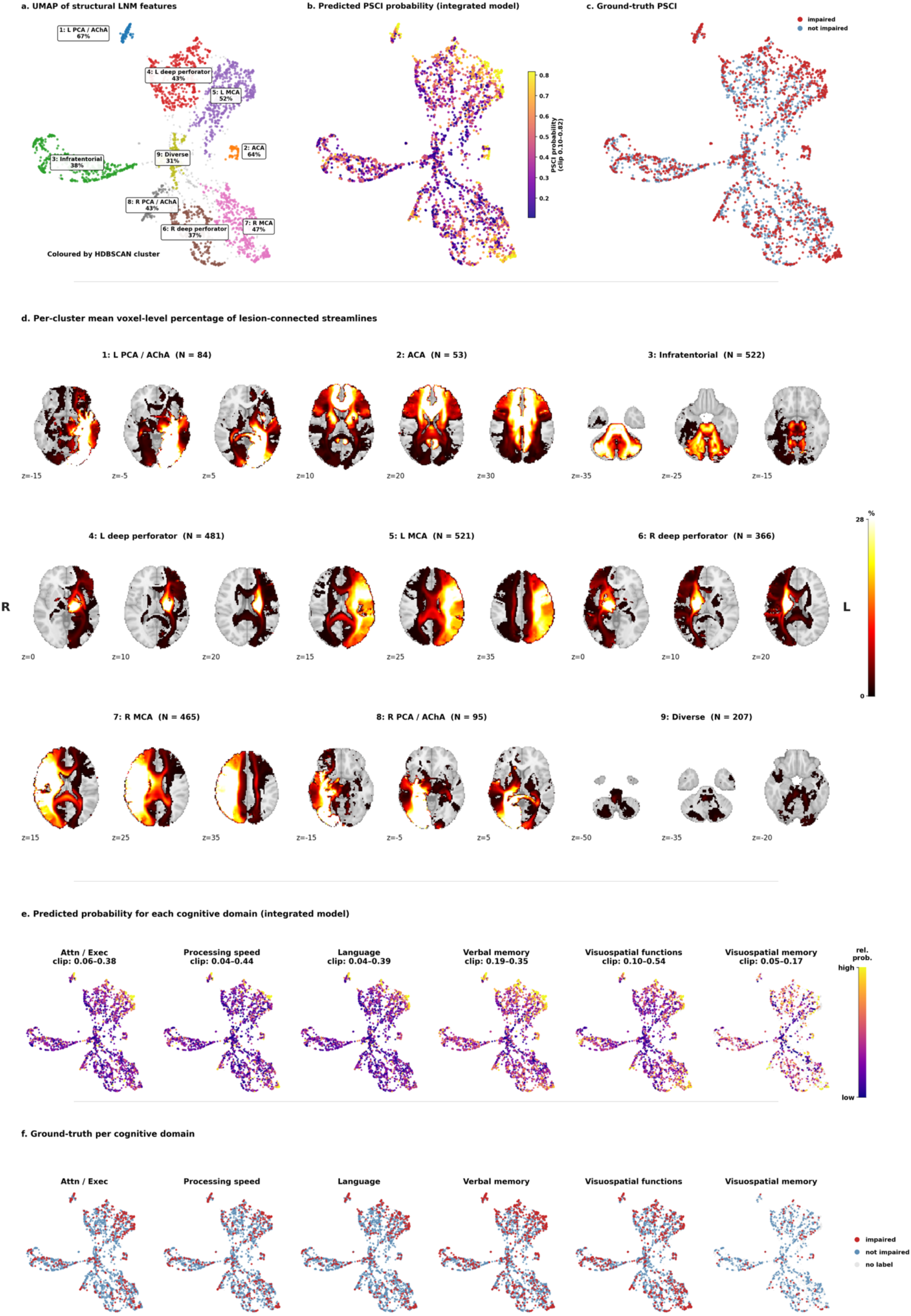
Data-driven patient stratification by structural lesion network mapping. **(a)** UMAP embedding of parcel-level structural LNM scores of acute infarcts, coloured by HDBSCAN cluster assignment. Nine clusters emerged, each labelled by its dominant vascular territory and observed any-domain PSCI prevalence. Of note, HDBSCAN does not necessarily assign every subject to a cluster; subjects not assigned to any cluster were labelled as noise and are displayed in lightgrey. **(b)** Same UMAP embedding coloured by the integrated model’s predicted any-domain PSCI probability, showing a gradient from low-risk (infratentorial, diverse) to high-risk (left PCA/AChA, ACA, left MCA) clusters. **(c)** Same embedding coloured by ground-truth any-domain PSCI status, confirming that the predicted probability gradient aligns with observed impairment rates. **(d)** Per-cluster mean structural lesion connectivity maps displayed at representative axial slices, illustrating the anatomical distribution of lesion-connected areas that defines each cluster. The colour scale indicates percentage of streamlines connected to the lesion. **(e)** Per-domain predicted impairment probability from the integrated model projected onto the UMAP embedding, with domain-specific colour scales clipped to the observed probability range. Predicted risk varied across the latent lesion connectivity landscape in a domain-specific manner — for example, left MCA and left deep perforator clusters show elevated predicted probabilities for impairment of attention / executive function, language and verbal memory, whereas right-hemispheric clusters show higher probabilities for visuospatial impairment. **(f)** Ground-truth impairment labels per cognitive domain projected onto the same embedding. Faded points indicate patients without available assessment for that domain. Embedding and clustering analyses of lesion location and functional LNM scores are provided in the *supplementary materials*. ACA = anterior cerebral artery; AChA = anterior choroidal artery; Attn/Exec. = attention/executive function; HDBSCAN = hierarchical density-based spatial clustering of applications with noise; L = left; MCA = middle cerebral artery; PCA = posterior cerebral artery; PSCI = post-stroke cognitive impairment; R = right; UMAP = uniform manifold approximation and projection.

### Pre-existing cerebrovascular burden further improves prediction of post-stroke cognitive impairment

Segmentations of pre-existing cerebrovascular lesions were available in a subset of patients (n = 1,973). We therefore evaluated pre-existing injury as an extension of the integrated model, compared with the acute infarct model fitted in the same subset. Adding pre-existing cerebrovascular injury to the integrated model further improved prediction (*figure 5a*). WMH topography and connectivity improved performance for five of seven cognitive outcomes, with the largest gains for processing speed (ΔAUPRC = +0.022) and attention/executive function (+0.019). Scores derived from the composite pre-existing lesion mask – combining WMH, old infarcts and lacunes per subject – improved performance for six of seven outcomes and were comparable in magnitude: larger than WMH alone for any-domain PSCI, language and visuospatial memory, and smaller for attention/executive function, processing speed and visuospatial function. Neither pre-existing lesion feature set improved prediction of verbal memory. Absolute performance of the baseline, integrated and extended models in this subset is shown in *supplementary figure S22*. Acute and pre-existing structural lesion connectivity were related to impairment in a graded manner. Any-domain PSCI prevalence increased across tertiles of both acute infarct and WMH structural lesion connectivity burden, and was highest when both were in the upper tertile (*figure 5b*). The same gradient was evident in all cognitive domains, though only weakly for visuospatial memory (*figure 5c*). Representing WMH by their topography and connectivity outperformed WMH volume alone, significantly improving prediction for three of seven outcomes (median ΔAUPRC = +0.04, up to +0.12 for any-domain PSCI; *figure 5d*).

**Figure 5.**
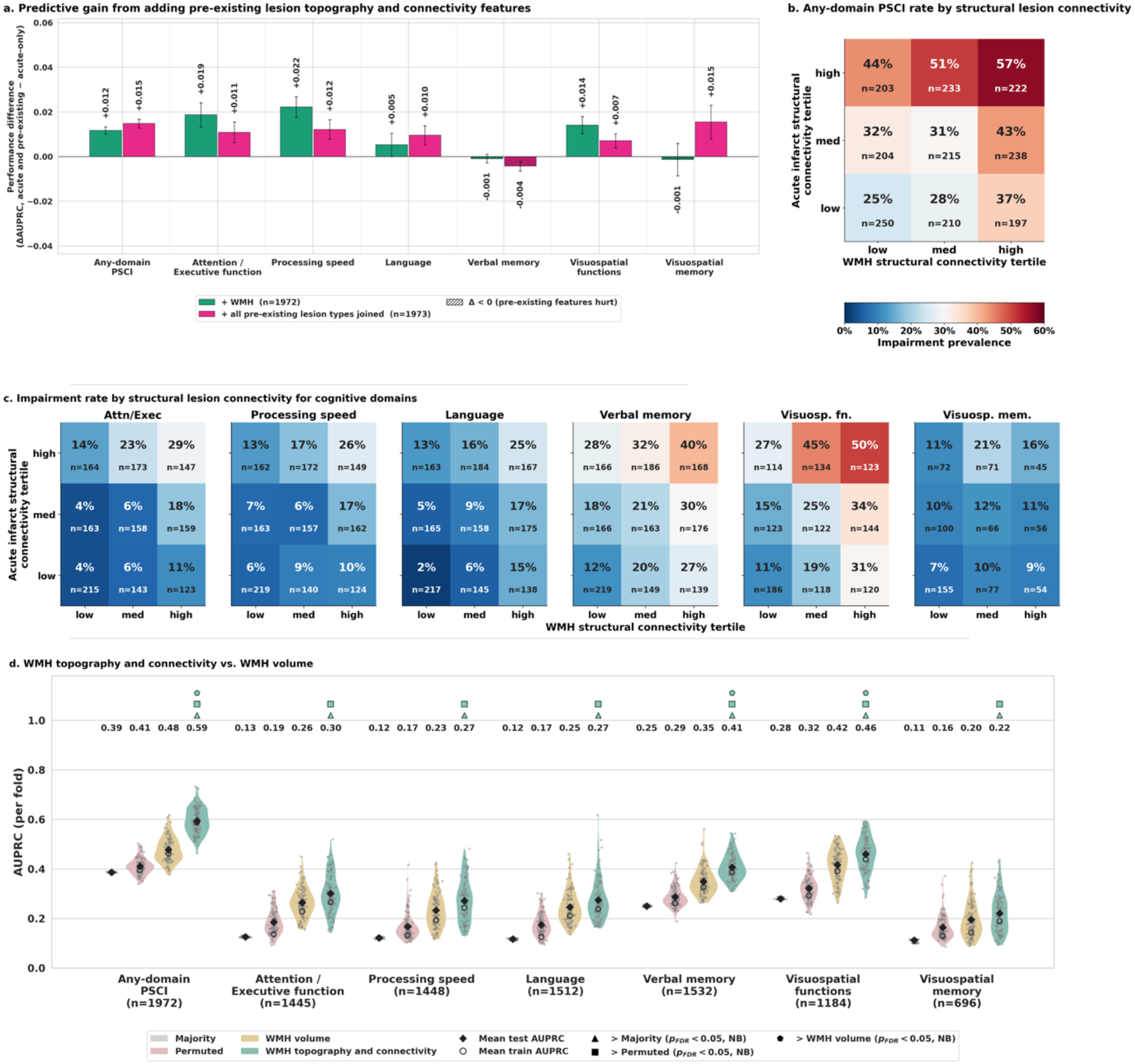
Predictive gain from adding pre-existing lesion features. **(a)** Predictive performance gain (ΔAUPRC) over the integrated acute infarct model from adding pre-existing lesion features. Pre-existing lesions were characterised using the same topography and connectivity representations as the acute infarct – i.e., lesion location, functional LNM, structural LNM – and entered the stacked ensemble as additional base learners. Two pre-existing feature sets are shown: WMH alone (green) and all pre-existing lesion types joined (magenta; composite: old infarcts, lacunes, and WMH combined). Hatched bars indicate targets where adding pre-existing lesion features reduced performance (Δ < 0). Adding WMH topography and connectivity yielded a modest AUPRC gain across five of the seven cognitive targets, whereas the pre-existing lesion union model showed similar improvements. For visuospatial memory prediction adding WMH features added no benefit, while adding composite features improved performance. **(b)** Observed any-domain PSCI prevalence stratified by tertiles of structural lesion connectivity (rows) and structural WMH connectivity (columns), showing that impairment risk increases along both axes and is highest when both lesion connectivity burdens are in the upper tertile. **(c)** The same tertile cross-tabulation for each of the six cognitive domains. Impairment rates increase along both axes – acute infarct and WMH connectivity burden – in every domain. **(d)** Predictive performance of models informed by WMH volume alone against WMH topography and connectivity. Models based on WMH topography and connectivity performed significantly better for three of seven cognitive targets. AUPRC = area under the precision–recall curve; WMH = white matter hyperintensities of presumed vascular origin; NB = Nadeau–Bengio; FDR = false discovery rate.

## Discussion

The cognitive consequences of focal brain injury should be considered from several complementary perspectives, from the site and extent of the lesion to the distributed brain systems it affects, to the pre-existing vulnerability of the brain in which it occurs. We showed that, in 3,034 patients from 13 international acute ischaemic stroke cohorts, modelling these aspects of the lesion–cognition relationship together predicted domain-specific post-stroke cognitive impairment: demographic and clinical characteristics with infarct volume predicted impairment well above chance; acute infarct topography and connectivity, structural lesion connectivity above all, significantly improved prediction for six of seven cognitive outcomes; and pre-existing cerebrovascular injury improved it further. The gains from lesion anatomy transferred to independent cohorts, most consistently for any-domain PSCI. The results highlighted left anterior thalamic connectivity as a shared predictor across most domains, alongside domain-specific anatomical profiles. Together, these findings support a lesion-in-context framework for understanding how the cognitive consequences of stroke emerge from the interaction between the lesion, the brain systems it affects, and patient-level factors.

### Structural lesion connectivity carried most of the predictive information beyond baseline

Removing structural LNM from the integrated model reduced predictive performance across all seven cognitive targets, while removing lesion location or functional LNM had little effect. In addition, structural LNM achieved the highest performance of the three feature sets when each was evaluated separately. These results should be interpreted in the context of the statistical dependence among the three lesion representations: all are deterministic transformations of the same binary infarct mask. Thus, the small penalty for removing lesion location or functional LNM likely reflects redundancy with structural LNM. For prediction, combining baseline information with structural LNM may therefore suffice. This interpretation is consistent with previous work showing that structural lesion connectivity provides greater predictive information than measures of immediate lesion location, while functional lesion connectivity offers comparatively limited additional predictive value.^7–9^ We extend this evidence to a large multi-centre setting.

More broadly, these findings suggest that enriching information about lesion location with network-level biological context can improve outcome prediction after acute brain injury. However, they should not be interpreted as establishing structural lesion connectivity as a biologically superior representation across all applications. Functional LNM, for instance, has proven valuable for localising lesion deficits, especially in rare syndromes with lesions in heterogeneous locations, and for identifying therapeutic stimulation targets.^16–18^ For predicting stroke outcomes, functional LNM may be less suited because its scores are more symmetric between hemispheres than lesion location or structural LNM scores, obscuring effects on lateralised cognitive functions, and because BOLD signal quality is lower in the subcortical and white matter regions that a lot of infarcts affect. Which representation is most valuable therefore depends on the question at hand, and understanding the information each captures remains an important challenge.^7,19–22^

### Sources and limits of the observed predictive benefit

Several analyses indicate that the predictive benefit reflects a reproducible but bounded signal. Performance increased with training set size and plateaued before reaching the full sample, while the advantage of the integrated model was present even in relatively small training samples. Thus, large samples were important for obtaining stable performance estimates, but the plateau indicates a ceiling that more data alone would not raise. This observation is consistent with evidence that reproducible brain–behaviour associations are generally small and require large samples to be estimated reliably.^23^ The broadly similar performance of ridge, XGBoost and TabPFN further suggests that the available signal is predominantly accessible to relatively simple models, with only modest evidence for additional nonlinear structure. Infarct size provides a complementary constraint: lesion topography and connectivity added little beyond the baseline for small infarcts and increasingly more for medium and large ones. Two explanations are plausible. Larger infarcts disrupt distributed systems more extensively, so that more of their cognitive consequence can be inferred from their anatomy, whereas after small infarcts variability may reside largely in other determinants. In addition, tracts and regions can lie millimetres from their position in normative atlases and connectomes, enough to decide whether a small infarct affects them. Because most infarcts in our sample were small, the pooled gain from lesion anatomy likely underestimates its value in more severe strokes, and performance figures should be read as conditional on case mix.

### Predictions relied on a shared thalamic core and domain-specific anatomy

Structural lesion connectivity of the left anterior thalamus carried a stable positive coefficient for every target and ranked among the most strongly weighted regions for any-domain PSCI, processing speed, language, verbal memory and visuospatial function; the adjacent left anterior thalamic radiation did so for any-domain PSCI, language and verbal memory. This convergence is consistent with the established role of thalamo-cortical and anterior thalamic–hippocampal circuitry in executive function, language and memory,^24–27^ and aligns with previous work identifying the left thalamus and its connected circuitry as a strategic substrate of vascular cognitive impairment, including post-stroke cognitive impairment.^28–31^ Beyond this shared core, most strongly weighted regions were specific to a single target. Domain-specific profiles consistent with prior literature included the left frontal aslant tract for verbal memory;^32^ the fornix and the right precuneus/posterior cingulate cortex for visuospatial memory;^33,34^ and dorsal attention and frontoparietal control networks for attention and executive function.^35,36^ Together with the spread of impairment across domains in our sample, this domain-specific anatomy underlines that PSCI is fully captured only by multi-domain assessment, and that differences in domain coverage may contribute to the variation in reported PSCI rates and lesion–cognition associations.

The lateralisation of several findings warrants some caution. The predominantly left-sided profiles for language, verbal memory and executive functions are neurobiologically plausible, but some nominally non-verbal neuropsychological measures also depend on verbal mediation. Consequently, left-lateralised predictive profiles should not necessarily be interpreted as purely domain-specific anatomical substrates. Similarly, the somatomotor contribution to different domains may partly reflect the graphomotor demands of these tests such as figure copying and timed visuomotor measures. As most patients were right-handed, the lateralised profiles may not extend to left-handed patients. These considerations favour interpreting the learned profiles as predictive anatomical signatures of the assessed cognitive phenotype, rather than as isolated causal substrates of individual cognitive functions.

Cerebellar regions also contributed to several domain-specific profiles. The cerebellum has established roles in executive, linguistic, visuospatial and affective functions, but the extent to which cerebellar involvement specifically contributes to PSCI remains less certain, particularly in heterogeneous cohorts with largely supratentorial stroke and potential crossed cerebellar diaschisis.^37^ The present findings therefore support cerebellar involvement as a hypothesis for further study.

### Structural lesion connectivity organised patients into recognisable vascular profiles

The coefficient profiles describe which connections contributed to prediction across the sample, but individual patients exhibit different combinations of lesion-related connectivity abnormalities. Grouping patients by these combinations yielded clusters that corresponded to the major vascular territories, consistent with disconnectome morphospace and generative lesion pattern approaches showing that lesion-derived features preserve the vascular organisation of stroke.^9,38^ Cohort membership and demographic characteristics were distributed diffusely across the embedding, suggesting that the grouping primarily reflected lesion anatomy.

The clusters differed in observed impairment prevalence, and predicted probabilities followed the same gradient. Notably, the ordering of risk did not follow territory size: infarcts of the left PCA/AChA and ACA territories, although the smallest clusters, carried higher prevalence than MCA infarcts, and their connectivity profiles centred on thalamic, medial temporal and midline frontal circuitry (*figure 4d*), converging with the thalamic core identified in the coefficient profiles and with strategic infarct locations reported previously.^28–31^ Prevalence was also consistently higher for left-than right-sided clusters of each paired territory, mirroring the lateralisation of the coefficient profiles.

### The acute lesion is one determinant among several

The baseline model already captured considerable variation in cognitive outcome, with demographic, clinical and infarct characteristics alone performing close to the integrated model. Its strongest contributors fall into two kinds: NIHSS and infarct volume reflect the severity and extent of the stroke, whereas cohort and time from stroke to cognitive assessment reflect the context of measurement, including recruitment, test batteries, normative data and recovery stage. Age and education, although established determinants of post-stroke cognition,^12^ contributed less, likely because impairment was defined against normative data that already account for them. Detailed lesion anatomy therefore added information beyond that already carried by these clinical and demographic variables.

### Pre-existing cerebrovascular injury provided a further, complementary dimension

Adding WMH topography and connectivity to the integrated model improved prediction for five of seven outcomes, while the composite representation incorporating WMH, old infarcts and lacunes improved prediction for six of seven outcomes. WMH added most for processing speed and attention/executive function, the domains classically associated with small vessel disease.^39^ Moreover, impairment increased progressively with both acute and pre-existing structural lesion connectivity burden and was highest when both were high. In addition, representing WMH by their topography and connectivity added predictive information beyond their volume, indicating that, as for the acute infarct, the cognitive impact of pre-existing lesions depends on where they lie and what they connect to, beyond how much tissue is affected. This is consistent with evidence from both memory clinic and stroke populations linking WMH distribution and connectivity to cognitive impairment.^11,29,40^ Together, these findings suggest that the limits of prediction reflect not only the complexity of the acutely available signal, but also the information about brain and patient state that is not captured by imaging of the acute infarct alone.

### The predictive benefit extended to independent cohorts

In leave-one-cohort-out cross-validation, the integrated model improved on the baseline for three of seven outcomes (four based on AUROC), and its benefit for any-domain PSCI was present in each of the 13 held-out cohorts. The benefit thus held across cohorts that differ in patient selection, imaging protocols, cognitive assessment and outcome prevalence,^15^ even though these differences lowered absolute performance relative to pooled cross-validation; the high contribution of cohort membership to the baseline model points to the same heterogeneity. This gap indicates that even data at this scale are not yet sufficient for reliable transfer across clinical settings. Closing it will require larger and more diverse datasets from which models can learn relationships robust to these sources of variation.

### From lesion anatomy to lesion context

Taken together, our findings indicate that the cognitive consequences of stroke cannot be read from the anatomy of the acute infarct alone: the lesion provides the starting point, but its impact depends on the circuitry it affects, the pre-existing state of the brain, and the characteristics of the individual.^4^ Jointly modelling these dimensions yielded consistent out-of-sample predictive improvements across cognitive domains that transferred to independent cohorts. Accordingly, an informative yet parsimonious model of post-stroke cognition would combine stroke severity, infarct volume and time to cognitive assessment with the structural connectivity of the acute infarct and the topography and connectivity of pre-existing cerebrovascular injury. Nevertheless, absolute performance remained modest. This is consistent with the broader observation that variables robustly associated with an outcome at the group level, including lesion–symptom associations, do not necessarily translate into accurate predictions for individual patients.^41^ Performance was further limited by the outcome itself: cognitive test scores have imperfect reliability and their harmonisation across cohorts will have introduced additional label noise that no predictor can capture. Determinants outside the lesion context captured here were not modelled, including pre-stroke cognitive reserve, concomitant neurodegeneration, genetic susceptibility, subacute cognitive function, and mood and alertness at assessment. Capturing these may be necessary for models accurate and transportable enough for trials and clinical care.

### Limitations

Cognitive impairment was represented as a binary label at a single time point within one year of stroke, simplifying both severity and longitudinal evolution. Information on premorbid cognitive ability and pre-stroke cognitive decline was not available. Both influence stroke risk, pre-existing vascular injury and post-stroke cognition, and education – which we included in our models – is only a partial proxy for them.^42,43^ Similarly, imaging features were restricted to vascular lesions on structural MRI and CT; brain morphometric markers capturing atrophy, and diffusion-based measures of white matter microstructure, were not available across cohorts and may add prognostic information. The inclusion of selected patient samples in several cohorts may limit generalisability to the broader clinical population. In particular, the relatively low median NIHSS and infarct volume indicate that our cohort predominantly comprised patients with mild strokes, and the observed associations may differ in patients with greater stroke severity or larger infarcts. Although cognitive measures were harmonised across cohorts and lesion masks were processed in a common template space, some residual heterogeneity from retrospective pooling of independently acquired cohorts, e.g., from differences in acquisition protocols or cognitive test administration, likely remains.^6,44^ Additionally, with most patients being of Asian and European ancestry, the generalisability of our findings to other ethnicities remains to be established.

## Conclusion

Across 3,034 patients with ischaemic stroke from 13 international cohorts, an informative model of cognitive outcome combined the size and structural connectivity of the acute infarct, the topography and connectivity of pre-existing cerebrovascular injury, and demographic and clinical characteristics, with gains that extended to independent cohorts. For future research, these findings support a lesion-in-context approach: understanding the cognitive consequences of focal damage requires considering not only where a lesion occurs, but also the distributed brain systems it affects and the broader brain and patient context that may shape its effects. Extending this approach to other lesion populations may help refine how lesion-based evidence is used to infer the neural systems underlying cognition.

## Methods

### Study population

We analysed previously harmonised, clinical and imaging data from 3,034 patients with acute ischaemic stroke enrolled across 13 Asian and European cohorts of the Meta VCI Map consortium (*figure 1a*). Meta VCI map is a multi-site collaboration for conducting meta- and mega-analyses of strategic lesion topography in vascular cognitive impairment.^44^ The stroke cohorts included in this study comprise Bundang VCI (n = 753, Korea), CASPER (n = 104, Netherlands), COAST (n = 74, Singapore), CODECS (n = 27, Netherlands), CROMIS-2 (n = 97, United Kingdom), CU-STRIDE (n = 410, Hong Kong), DEDEMAS (n = 87, Germany), GRECogVASC (n = 313, France), Hallym VCI (n = 641, Korea), MSS-2 (n = 100, United Kingdom), PROCRAS (n = 177, Netherlands), STROKDEM (n = 138, France), USCOG (n = 113, Netherlands). Compared with the original dataset analysed by Weaver et al.,^6^ the GRECogVASC data have been updated and DEDEMAS has been added, so that cohort sizes and descriptive statistics differ from the previous publication. All cohorts include patients with acute ischaemic stroke; available brain CT or MRI (T1-weighted, T2-weighted, fluid-attenuated inversion recovery [FLAIR], or diffusion-weighted imaging [DWI]) showing the symptomatic infarcts; available infarct segmentations; and a study protocol that includes cognitive assessment within 1 year of the index stroke.^6^ Patients without data for age, sex, level of education, or a cognitive assessment within 1 year of the index stroke, consisting of either a multidomain neuropsychological assessment with data on ≥3 cognitive domains or the Montreal Cognitive Assessment, were excluded.^6^

### Ethics approval

All cohorts received the requisite ethical and institutional approval_in accordance with local regulations, which included informed consent, to allow data acquisition and sharing.^6^

### Cognitive assessments

Detailed harmonisation procedures, including the assignment of individual tests to cognitive domains, have been reported previously.^6^ Neuropsychological tests administered within 1 year after stroke were norm-referenced against local normative data. Performance below the fifth percentile was considered impaired. Tests were assigned to one of six cognitive domains: attention/executive function, processing speed, language, verbal memory, visuospatial function, and visuospatial memory. For each individual, a cognitive domain was classified as impaired when more than 50% of the available tests within that domain were impaired. Any-domain PSCI was defined as impairment in at least one cognitive domain. In three cohorts (CROMIS-2, CU-STRIDE and MSS-2), only the MoCA was available; in these cohorts, any-domain PSCI was defined as a total MoCA score below the fifth percentile of age- and education-corrected normative data, as previously described.^6^ These cohorts therefore contributed to the any-domain PSCI outcome only, and all domain-specific outcomes are based exclusively on multidomain neuropsychological assessment.

### Lesion segmentations

Lesion segmentations were provided by the participating centres or, where unavailable, performed centrally at UMC Utrecht. Lesion masks for acute and prior infarcts were obtained based on CT and MRI scans, depending on their availability. WMH and lacunes were segmented based on MRI. Segmentation masks were spatially normalised to the MNI152NLin6Asym template.^45^ To ensure registration quality, the normalised lesion masks were visually inspected and patients with failed registrations were excluded. Further details on quality control and cohort-specific segmentation and registration procedures were reported previously.^6^

### Feature extraction

For each acute infarct, we calculated three scores for each of 190 prespecified brain regions (*figure 1b*): lesion location scores, functional LNM scores, and structural LNM scores. Together, these scores captured complementary aspects of lesion anatomy and its effects on the brain. Lesion location scores quantified direct tissue damage as the percentage of each region’s volume intersected by the lesion mask. Functional LNM scores quantified the functional connectivity of each region to the lesion as the mean t-statistic within that region of a lesion-seeded connectivity map derived from a normative resting-state connectome (GSP1000).^16,46,47^ Structural LNM scores quantified the structural connectivity of each region to the lesion as the proportion of streamlines associated with that region that passed through the lesion, based on a normative tractogram (dTOR985).^48^ Further methodological details on the derivation of the functional and structural LNM scores are provided in *supplementary text S1*.

The 190 brain regions were defined by a standardised parcellation: Schaefer cortex (n_ROIs_ = 100), Tian subcortex (n_ROIs_ = 16), Nettekoven cerebellum (n_ROIs_ = 32) and XTRACT white matter tracts (n_ROIs_ = 42).^49–52^ We additionally characterised pre-existing cerebrovascular lesions using the same three imaging representations. Segmentations of WMH, lacunes and old non-lacunar infarcts were available for 1,973 patients; WMH were present in 1,972, lacunes in 773 and old infarcts in 341. Feature sets were derived for two types of lesion masks. The first was the WMH mask alone (n_patients_ = 1,972). The second was a composite mask formed per patient as the voxel-wise union of all lesion types present (n_patients_ = 1,973); in patients with WMH only, the composite therefore equalled the WMH mask. Features were extracted from this single binary mask. This composite treats all pre-existing lesion types as equivalent and disregards differences in how they affect tissue; we adopted this simplification because old infarcts and lacunes were present in too few patients to model each lesion type separately.

The complete feature extraction workflow — from lesion mask to lesion location, functional and structural lesion network scores — is released alongside this work as part of Lacuna, an open-source toolbox (lacuna-py.readthedocs.io).^53^ Lacuna requires a binary lesion mask in standard space (MNI152NLin6Asym or MNI152NLin2009cAsym) as input and provides an easy-to-use interface to derive all features used here, enabling reproducible extraction across studies.

Prediction framework.

Our predictive modelling pipeline was designed to minimise overfitting, data leakage, and circularity in accordance with current best practices for machine learning neuroimaging.^54^ We first evaluated whether integrating complementary representations of lesion anatomy improves prediction of post-stroke cognitive impairment beyond conventional clinical information. To address this, we compared four models of increasing complexity: (1) a majority-class reference model; (2) a null model trained on permuted outcome labels; (3) a baseline model representing the information routinely available to the clinician, comprising demographic characteristics (age, sex, education level, cohort, ethnicity), clinical variables (NIHSS, handedness, prior stroke, prior transient ischaemic attack, interval stroke onset until cognitive assessment, interval stroke onset until MRI, obesity, arterial hypertension, hyperlipidaemia, diabetes mellitus, present smoking, past smoking), and lesion volume as a basic imaging measure; and (4) an integrated model combining the baseline variables with lesion location, functional LNM, and structural LNM scores. All models used identical preprocessing. Within each outer training fold, missing values in the demographic and clinical variables were replaced by the training fold median, and all model inputs were z-scored using training fold means and standard deviations. Both transformations were estimated on the training fold only and applied unchanged to the held-out test fold to avoid data leakage.^55^

The integrated model combined the feature sets using a stacked ensemble. Ridge classifiers with internally cross-validated regularisation served as base learners: one trained on the baseline variables alone, and one for each imaging representation (lesion location, functional LNM, and structural LNM), with each imaging model incorporating the baseline variables. Within each outer training fold, base learner predictions were generated as out-of-fold predictions using 10-fold inner cross-validation. The resulting out-of-fold prediction matrix was standardised and combined by an L2-regularised logistic regression meta-learner. The base learners were subsequently refit on the full outer training fold to generate predictions for the held-out test fold. Because the meta-learner was trained exclusively on out-of-fold predictions, it learned to combine complementary information across representations without leaking information into the evaluation process.^55^

All models were trained and evaluated using identical fold assignments under pooled 10 × 10 repeated stratified cross-validation, enabling paired model comparisons. Hyperparameter optimisation was nested entirely within the training data at every stage: Ridge regularisation strengths were selected by leave-one-out cross-validation within each base learner, while the meta-learner’s regularisation parameter was selected by 10-fold cross-validation on the out-of-fold prediction matrix. Thus, imputation, input scaling, model fitting, stacking, and hyperparameter optimisation were all performed exclusively within the outer training folds, with the held-out test folds reserved solely for unbiased performance estimation. Predictive performance was quantified as the AUPRC.^56^ Chance-level AUPRC corresponds to the impairment prevalence of each cognitive target. Calibration of the integrated model was assessed from out-of-fold predicted probabilities using calibration-in-the-large and the calibration slope (*supplementary figure S3*). Models were compared using one-sided Nadeau–Bengio paired *t*-tests with Benjamini–Hochberg false-discovery-rate correction within each cognitive endpoint.^57^

### Sources and robustness of predictive performance

We performed four complementary analyses to better understand the determinants of model performance. First, ablation analyses quantified the unique contribution of each imaging representation by systematically removing lesion location, functional LNM, or structural LNM scores from the integrated model. Second, learning curves assessed how predictive accuracy changed as a function of training sample size.^58^ Third, we evaluated whether the findings depended on the prediction algorithm by comparing the integrated model based on the linear stacking framework with XGBoost and TabPFN under an otherwise identical evaluation framework (for details see *supplementary text S2*).^59,60^ Finally, we examined whether the benefit of detailed lesion characterisation depended on patient characteristics by repeating the analysis within strata of infarct volume (<5 mL, 5–30 mL and ≥30 mL), age (tertiles) and sex. Within each stratification, the stratifying variable was removed from the baseline variables.

### Cross-cohort generalisation

We then evaluated whether the predictive benefit of integrating complementary imaging representations extended to independent clinical cohorts. The integrated model was re-evaluated under leave-one-cohort-out cross-validation, in which each cohort served once as a held-out test set while the model was trained on the remaining cohorts. Imputation and scaling were re-estimated on the training cohorts at each split, so that no information from the held-out cohort informed preprocessing. The modelling framework was otherwise identical to the pooled analysis: the same feature sets, the same stacked ensemble of ridge base learners and logistic regression meta-learner, and the same nested hyperparameter optimisation and out-of-fold stacking procedure, all performed exclusively within the training cohorts. Two aspects differed: cohort membership instead of repeated stratified folds defined the train–test split, and the cohort indicator was removed from the baseline variables. To account for differences in impairment prevalence between cohorts, performance was expressed as normalised AUPRC ((AUPRC – prevalence)/(1 – prevalence)). The additional predictive value of the integrated model over the baseline was assessed across held-out cohorts using paired Wilcoxon signed-rank tests.

### Neuroanatomical interpretation

Next, we examined which anatomical profiles contributed to improvements in predictive performance. Because ablation analyses identified structural LNM as the most informative imaging feature set, we focused this analysis on the structural LNM base learner. Ridge coefficients were averaged across all cross-validation folds, identifying robust anatomical predictors across cortical, subcortical, cerebellar, and white matter regions.^13^ Because model coefficients reflect how information is combined across correlated regions, they indicate which regions the model relied on rather than the strength of each region’s association with impairment.^61^ For completeness, the same procedure was applied to the lesion location and functional LNM base learners, with the results presented in the *supplementary materials*.

Whereas the coefficient analysis describes the whole sample, individual patients differ in which combinations of connections their infarct implicates. To characterise this variation, we additionally grouped patients by the similarity of their structural lesion connectivity. Patient-level structural LNM scores were projected into a low-dimensional embedding using UMAP (n_neighbors_ = 30, min_dist_ = 0.1, n_components_ = 2, Euclidean metric).^62^ Clusters were identified by applying HDBSCAN to the two-dimensional embedding.^63^ Both steps were performed unsupervised, i.e., without reference to cognitive outcome. Cluster membership was then related to observed impairment prevalence and to the predicted probabilities of the integrated model.

### Contribution of pre-existing cerebrovascular injury

We assessed whether pre-existing cerebrovascular injury contributes predictive information beyond the acute infarct. To address this, we extended the integrated acute infarct model with the lesion location, functional LNM, and structural LNM scores derived from the two pre-existing lesion masks described above, yielding two extended models: one adding the WMH scores and one adding the scores derived from the composite pre-existing lesion mask. Improvements in predictive performance were quantified as changes in AUPRC for each cognitive outcome, relative to the acute infarct-only model fitted in the same patients. To determine whether the anatomical distribution of WMH provides information beyond overall lesion burden, we additionally compared a model based on the three types of WMH scores – lesion location, functional LNM, structural LNM – with one based on WMH volume alone.

### Sensitivity analyses

We tested whether predictive modelling results depended on four methodological choices in deriving lesion features. First, to test the influence of parcellation resolution, the integrated model was refit on lesion location, functional LNM and structural LNM scores based on two higher-resolution atlas sets, a mid-resolution set (Schaefer-400 cortex, Tian S2 subcortex, Nettekoven-32 cerebellum, XTRACT) and a high-resolution set (Schaefer-1000, Tian S4, Nettekoven-64, XTRACT). These models were compared with the core integrated model based on the 190-region reference parcellation. Second, because computing structural LNM scores from tractograms is resource-intensive, we tested whether sparser versions of dTOR985, which lower the computational cost of structural LNM, yield equivalent results. Structural LNM scores were recomputed from random subsamples of 10%, 25%, 50% and 75% of the dTOR985 streamlines and compared with the full tractogram. Third, because infarct volume was strongly right-skewed, we replaced raw infarct volume by its log-transformed value in all models to test whether its linear representation in the baseline affected the results. Fourth, because previous lesion-based prediction studies have used voxelwise lesion maps reduced by principal component analysis instead of region-wise scores, we compared both feature formats for each lesion representation.^7,10,64^ For lesion location, functional LNM and structural LNM, single-view models (baseline plus one representation) were fitted either on the 190 regional scores or on the voxelwise maps (2 mm) reduced to the principal components explaining 95% of their variance, with the decomposition estimated within each training fold. All variants were evaluated on identical folds within the pooled 10 × 10 repeated stratified cross-validation framework.

## Acknowledgments.

The authors wish to acknowledge all participants of the stroke group of the Meta VCI Map consortium. The Meta VCI Map stroke project involved all listed authors, as well as support from Jill Abrigo, Mélanie Barbay, Guido Cammà, Anna Dewenter, Martin Dichgans, Thomas Doeven, Marco Duering, Lieza G. Exalto, Rong Fang, Bibek Gyanwali, Olivia K.L. Hamilton, Irene M.C. Huenges Wajer, L. Jaap Kappelle, Beom Joon Kim, Peter J. Koudstaal, Gregory Kuchcinski, Hugo J. Kuijf, Anne-Marie Mendyk, Robert J. van Oostenbrugge, Lin Shi, Charlotte M. Verhagen, Frans R.J. Verhey, Martine J.E. van Zandvoort, and Lei Zhao. During the preparation of this manuscript, the authors used Claude (Anthropic) to improve language and readability of author-written text.

## Code availability

All analysis code is publicly available on GitHub (https://github.com/umcu-VCI-group/2026_petersen-lesion_in_context_psci). Lesion location, functional lesion network mapping and structural lesion network mapping features were computed with Lacuna.^53^ Analyses were performed in Python 3.11 using scikit-learn 1.8.0, XGBoost 3.2.0, TabPFN 6.4.1, NumPy 2.3.5, pandas 2.3.3, SciPy 1.15.3, joblib 1.5.3, nibabel 5.3.3, nilearn 0.13.0, umap-learn 0.5.12 and hdbscan 0.8.43; figures were produced with matplotlib 3.10.8, seaborn 0.13.2, adjustText 1.3.0, PyVista 0.47.3 and yabplot 0.4.0.

## Data availability

The Meta VCI Map consortium is dedicated to data sharing. The combined dataset of the present study, including deidentified participant data and lesion maps in MNI space, can be used for additional analyses on reasonable request. Any request requires approval of the Meta VCI Map project group, consisting of representatives of all participating cohorts. Participating cohort groups have shared their data with the consortium under the agreement that the source data remains property of the group that supplied the data. Source data are stored at the participating centres and can be requested from each group.

## Contributors

Each author has made a significant contribution to the manuscript and all authors read and approved its final version. We describe contributions to the paper using the CRediT contributor role taxonomy. Conceptualization: MP, GJB. Data curation: MP, NAW, JMB, FASdK, MC, GJB. Methodology: MP, KRP, MDF, AH, FLWVJS, SBE, GJB. Software: MP. Formal analysis: MP. Investigation: all authors. Resources: all authors. Visualization: MP. Writing – original draft: MP. Writing – review & editing: all authors. Supervision: KRP, SBE, GJB. Project administration: MP, GJB. Funding acquisition: MP, GJB. MP, NAW and GJB have accessed and verified the underlying data.

