## Supplementary materials for "Determinants of domain-specific cognitive impairment after stroke: a mega-analysis of 13 acute ischaemic stroke cohorts"

**This PDF file includes:**

Figs. S1 to S22
Texts S1 to S2
Table S1
References

### Table S1 | Characteristics of the study population by cohort.

| Characteristic | Pooled | CASPER | COAST | CODECS | CROMIS-2 | DEDEMAS | PROCRAS | GRECogVASC | Hallym VCI | STROKDEM | MSS-2 | Bundang VCI | CU-STRIDE | USCOG |
|---|---|---|---|---|---|---|---|---|---|---|---|---|---|---|
| **N** | **3,034** | **104** | **74** | **27** | **97** | **87** | **177** | **313** | **641** | **138** | **100** | **753** | **410** | **113** |
| **Demographics** | | | | | | | | | | | | | | |
| Age (years) | 68.0 [59.0–76.0] 66.8 ± 11.6 | 66.2 [56.3–71.6] 64.1 ± 10.8 | 58.5 [51.0–64.0] 58.4 ± 10.5 | 54.0 [51.0–72.0] 59.2 ± 16.5 | 75.0 [67.0–79.0] 73.7 ± 9.2 | 72.0 [64.5–76.0] 70.1 ± 8.3 | 70.0 [65.0–76.0] 69.6 ± 9.4 | 65.0 [55.0–74.0] 63.7 ± 10.7 | 66.0 [56.0–74.0] 65.1 ± 11.9 | 66.0 [57.0–74.8] 64.9 ± 12.1 | 66.0 [58.5–75.0] 65.7 ± 11.6 | 71.0 [64.0–77.0] 69.8 ± 10.8 | 69.0 [62.0–77.0] 68.6 ± 10.4 | 61.0 [51.0–71.0] 60.0 ± 14.9 |
| **Sex** | | | | | | | | | | | | | | |
| Female | 1,184 (39.0%) | 27 (26.0%) | 21 (28.4%) | 12 (44.4%) | 43 (44.3%) | 27 (31.0%) | 58 (32.8%) | 121 (38.7%) | 268 (41.8%) | 53 (38.4%) | 38 (38.0%) | 306 (40.6%) | 163 (39.8%) | 47 (41.6%) |
| Male | 1,850 (61.0%) | 77 (74.0%) | 53 (71.6%) | 15 (55.6%) | 54 (55.7%) | 60 (69.0%) | 119 (67.2%) | 192 (61.3%) | 373 (58.2%) | 85 (61.6%) | 62 (62.0%) | 447 (59.4%) | 247 (60.2%) | 66 (58.4%) |
| **Ethnicity** | | | | | | | | | | | | | | |
| Korean | 1,394 (46.0%) | 0 (0.0%) | 0 (0.0%) | 0 (0.0%) | 0 (0.0%) | 0 (0.0%) | 0 (0.0%) | 0 (0.0%) | 641 (100.0%) | 0 (0.0%) | 0 (0.0%) | 753 (100.0%) | 0 (0.0%) | 0 (0.0%) |
| Malay | 16 (0.5%) | 0 (0.0%) | 16 (21.6%) | 0 (0.0%) | 0 (0.0%) | 0 (0.0%) | 0 (0.0%) | 0 (0.0%) | 0 (0.0%) | 0 (0.0%) | 0 (0.0%) | 0 (0.0%) | 0 (0.0%) | 0 (0.0%) |
| Indian | 6 (0.2%) | 0 (0.0%) | 6 (8.1%) | 0 (0.0%) | 0 (0.0%) | 0 (0.0%) | 0 (0.0%) | 0 (0.0%) | 0 (0.0%) | 0 (0.0%) | 0 (0.0%) | 0 (0.0%) | 0 (0.0%) | 0 (0.0%) |
| Chinese | 462 (15.2%) | 0 (0.0%) | 52 (70.3%) | 0 (0.0%) | 0 (0.0%) | 0 (0.0%) | 0 (0.0%) | 0 (0.0%) | 0 (0.0%) | 0 (0.0%) | 0 (0.0%) | 0 (0.0%) | 410 (100.0%) | 0 (0.0%) |
| European | 1,154 (38.1%) | 104 (100.0%) | 0 (0.0%) | 27 (100.0%) | 95 (100.0%) | 87 (100.0%) | 177 (100.0%) | 313 (100.0%) | 0 (0.0%) | 138 (100.0%) | 100 (100.0%) | 0 (0.0%) | 0 (0.0%) | 113 (100.0%) |
| *Missing* | *2 (0.1%)* | *0* | *0* | *0* | *2 (2.1%)* | *0* | *0* | *0* | *0* | *0* | *0* | *0* | *0* | *0* |
| **Education (STROKOG)** | | | | | | | | | | | | | | |
| Level 1 | 1,757 (57.9%) | 42 (40.4%) | 52 (70.3%) | 8 (29.6%) | 6 (6.2%) | 55 (63.2%) | 79 (44.6%) | 232 (74.1%) | 351 (54.8%) | 86 (62.3%) | 19 (19.0%) | 402 (53.4%) | 381 (92.9%) | 44 (38.9%) |
| Level 2 | 614 (20.2%) | 19 (18.3%) | 14 (18.9%) | 4 (14.8%) | 78 (80.4%) | 7 (8.0%) | 53 (29.9%) | 29 (9.3%) | 154 (24.0%) | 16 (11.6%) | 62 (62.0%) | 146 (19.4%) | 5 (1.2%) | 27 (23.9%) |
| Level 3 | 257 (8.5%) | 35 (33.7%) | 7 (9.5%) | 5 (18.5%) | 0 (0.0%) | 24 (27.6%) | 39 (22.0%) | 34 (10.9%) | 31 (4.8%) | 11 (8.0%) | 0 (0.0%) | 37 (4.9%) | 8 (2.0%) | 26 (23.0%) |
| Level 4 | 406 (13.4%) | 8 (7.7%) | 1 (1.4%) | 10 (37.0%) | 13 (13.4%) | 1 (1.1%) | 6 (3.4%) | 18 (5.8%) | 105 (16.4%) | 25 (18.1%) | 19 (19.0%) | 168 (22.3%) | 16 (3.9%) | 16 (14.2%) |

| Characteristic | Pooled | CASPER | COAST | CODECS | CROMIS-2 | DEDEMAS | PROCRAS | GRECogVASC | Hallym VCI | STROKDEM | MSS-2 | Bundang VCI | CU-STRIDE | USCOG |
|---|---|---|---|---|---|---|---|---|---|---|---|---|---|---|
| **Handedness** | | | | | | | | | | | | | | |
| Left | 109 (3.9%) | 6 (5.8%) | — | 5 (20.0%) | 8 (8.4%) | — | 6 (3.4%) | 31 (9.9%) | 6 (1.0%) | 10 (13.5%) | 11 (11.0%) | 7 (0.9%) | 7 (1.8%) | 12 (10.7%) |
| Right | 2,617 (94.6%) | 95 (91.3%) | — | 20 (80.0%) | 87 (91.6%) | — | 167 (94.4%) | 282 (90.1%) | 609 (98.2%) | 64 (86.5%) | 89 (89.0%) | 726 (96.7%) | 380 (96.4%) | 98 (87.5%) |
| Ambidextrous | 39 (1.4%) | 3 (2.9%) | — | 0 (0.0%) | 0 (0.0%) | — | 4 (2.3%) | 0 (0.0%) | 5 (0.8%) | 0 (0.0%) | 0 (0.0%) | 18 (2.4%) | 7 (1.8%) | 2 (1.8%) |
| *Missing* | *269 (8.9%)* | *0* | *74 (100.0%)* | *2 (7.4%)* | *2 (2.1%)* | *87 (100.0%)* | *0* | *0* | *21 (3.3%)* | *64 (46.4%)* | *0* | *2 (0.3%)* | *16 (3.9%)* | *1 (0.9%)* |
| **Stroke and imaging characteristics** | | | | | | | | | | | | | | |
| NIHSS | 3.0 [1.0–5.0] 3.9 ± 4.5 | — | 3.0 [1.0–7.0] 4.5 ± 4.3 | 0.0 [0.0–0.0] 0.4 ± 0.8 | 3.0 [2.0–6.0] 5.4 ± 6.1 | 2.0 [1.0–4.0] 3.8 ± 4.7 | 3.0 [2.0–4.0] 3.9 ± 3.9 | 3.0 [1.0–6.0] 4.9 ± 5.3 | 2.0 [1.0–4.0] 3.2 ± 3.9 | 0.0 [0.0–1.0] 0.7 ± 1.3 | 1.0 [0.0–2.0] 1.1 ± 1.2 | 3.0 [2.0–5.0] 4.5 ± 4.8 | 4.0 [2.0–6.0] 4.9 ± 4.6 | — |
| *Missing* | *304 (10.0%)* | *104 (100.0%)* | *0* | *0* | *40 (41.2%)* | *8 (9.2%)* | *0* | *0* | *33 (5.1%)* | *2 (1.4%)* | *0* | *0* | *4 (1.0%)* | *113 (100.0%)* |
| Prior stroke | 336 (11.1%) | 5 (4.8%) | 10 (13.5%) | 0 (0.0%) | 6 (6.2%) | 9 (10.3%) | 23 (13.0%) | 22 (7.0%) | 85 (13.5%) | 12 (8.7%) | 10 (10.0%) | 104 (13.8%) | 50 (12.2%) | 0 (0.0%) |
| *Missing* | *13 (0.4%)* | *0* | *0* | *0* | *1 (1.0%)* | *0* | *0* | *0* | *12 (1.9%)* | *0* | *0* | *0* | *0* | *0* |
| Prior TIA | 97 (3.8%) | 2 (1.9%) | — | 0 (0.0%) | 9 (9.5%) | 3 (3.4%) | 23 (13.0%) | — | 5 (0.9%) | 7 (5.1%) | 8 (8.0%) | 15 (2.0%) | 6 (1.5%) | 19 (17.3%) |
| *Missing* | *452 (14.9%)* | *0* | *74 (100.0%)* | *0* | *2 (2.1%)* | *0* | *0* | *313 (100.0%)* | *60 (9.4%)* | *0* | *0* | *0* | *0* | *3 (2.7%)* |
| Onset to NPA (days) | 105 [74–170] 117 ± 79 | 87 [81–98] 90 ± 15 | 120 [105–151] 136 ± 42 | 90 [90–90] 90 ± 0 | 4 [2–9] 7 ± 7 | — | 35 [29–40] 35 ± 10 | 178 [161–186] 177 ± 19 | 98 [90–105] 104 ± 30 | 189 [178–199] 201 ± 48 | 142 [53–383] 213 ± 166 | 104 [10–170] 109 ± 93 | 154 [129–176] 154 ± 37 | 6 [4–9] 7 ± 5 |
| *Missing* | *94 (3.1%)* | *0* | *0* | *0* | *1 (1.0%)* | *87 (100.0%)* | *0* | *0* | *3 (0.5%)* | *0* | *3 (3.0%)* | *0* | *0* | *0* |
| Onset to MRI (days) | 4 [1–9] 28 ± 59 | 87 [81–99] 88 ± 22 | 2 [1–4] 16 ± 74 | 34 [5–97] 61 ± 67 | 5 [3–9] 6 ± 5 | 3 [2–5] 4 ± 2 | 33 [27–40] 34 ± 9 | 178 [161–186] 177 ± 19 | 1 [1–2] 2 ± 3 | 3 [3–3] 3 ± 1 | 4 [3–9] 7 ± 8 | 5 [4–6] 5 ± 3 | 1 [0–2] 1 ± 1 | 5 [3–8] 35 ± 96 |
| *Missing* | *5 (0.2%)* | *0* | *0* | *1 (3.7%)* | *0* | *0* | *0* | *0* | *3 (0.5%)* | *0* | *0* | *1 (0.1%)* | *0* | *0* |
| Total infarct volume (mL) | 2.9 [1.0–14.7] 15.7 ± 36.8 | 3.4 [0.9–13.1] 11.2 ± 19.2 | 6.8 [2.0–30.6] 26.9 ± 49.5 | 10.3 [1.2–24.6] 19.4 ± 23.7 | 4.5 [1.5–15.9] 13.9 ± 23.9 | 1.9 [0.6–11.1] 9.9 ± 21.9 | 4.4 [1.3–21.0] 15.2 ± 24.7 | 1.9 [0.4–11.1] 14.0 ± 33.4 | 2.0 [0.9–11.2] 10.8 ± 22.7 | 1.7 [0.6–8.3] 7.7 ± 17.6 | 2.6 [1.2–10.7] 9.7 ± 20.9 | 3.8 [1.2–16.4] 21.9 ± 53.6 | 2.3 [0.9–12.9] 11.8 ± 28.4 | 19.6 [3.5–51.4] 38.0 ± 50.6 |

| Characteristic | Pooled | CASPER | COAST | CODECS | CROMIS-2 | DEDEMAS | PROCRAS | GRECogVASC | Hallym VCI | STROKDEM | MSS-2 | Bundang VCI | CU-STRIDE | USCOG |
|---|---|---|---|---|---|---|---|---|---|---|---|---|---|---|
| **Vascular risk factors** | | | | | | | | | | | | | | |
| Obesity | 857 (36.7%) | — | — | 4 (14.8%) | — | — | 46 (26.1%) | 214 (69.0%) | 219 (35.8%) | 35 (25.4%) | — | 250 (33.4%) | 89 (27.7%) | — |
| *Missing* | *702 (23.1%)* | *104 (100.0%)* | *74 (100.0%)* | *0* | *97 (100.0%)* | *87 (100.0%)* | *1 (0.6%)* | *3 (1.0%)* | *29 (4.5%)* | *0* | *100 (100.0%)* | *5 (0.7%)* | *89 (21.7%)* | *113 (100.0%)* |
| Smoking (current) | 682 (23.2%) | 18 (17.3%) | 22 (29.7%) | — | 11 (11.5%) | 46 (52.9%) | 38 (21.5%) | 75 (24.0%) | 187 (29.5%) | 31 (22.5%) | 29 (29.0%) | 158 (21.0%) | 50 (12.2%) | 17 (34.7%) |
| *Missing* | *100 (3.3%)* | *0* | *0* | *27 (100.0%)* | *1 (1.0%)* | *0* | *0* | *0* | *8 (1.2%)* | *0* | *0* | *0* | *0* | *64 (56.6%)* |
| Smoking (past) | 629 (23.1%) | 61 (58.7%) | 4 (5.4%) | 7 (25.9%) | 42 (43.8%) | — | 84 (47.5%) | 56 (17.9%) | 58 (9.4%) | — | 32 (32.0%) | 149 (19.8%) | 125 (30.5%) | 11 (22.4%) |
| *Missing* | *314 (10.3%)* | *0* | *0* | *0* | *1 (1.0%)* | *87 (100.0%)* | *0* | *0* | *24 (3.7%)* | *138 (100.0%)* | *0* | *0* | *0* | *64 (56.6%)* |
| Hypertension | 2,012 (67.7%) | 80 (76.9%) | 56 (75.7%) | 15 (55.6%) | 54 (56.2%) | 52 (59.8%) | 132 (74.6%) | 183 (58.5%) | 385 (60.2%) | 76 (55.1%) | 76 (76.0%) | 578 (76.8%) | 305 (74.4%) | 20 (38.5%) |
| *Missing* | *63 (2.1%)* | *0* | *0* | *0* | *1 (1.0%)* | *0* | *0* | *0* | *1 (0.2%)* | *0* | *0* | *0* | *0* | *61 (54.0%)* |
| Hyperlipidaemia | 1,362 (46.0%) | 88 (84.6%) | 59 (79.7%) | 9 (33.3%) | 48 (50.0%) | 30 (34.5%) | 167 (94.4%) | 136 (43.5%) | 246 (38.9%) | 60 (43.5%) | 64 (64.0%) | 201 (26.7%) | 242 (59.0%) | 12 (23.1%) |
| *Missing* | *70 (2.3%)* | *0* | *0* | *0* | *1 (1.0%)* | *0* | *0* | *0* | *8 (1.2%)* | *0* | *0* | *0* | *0* | *61 (54.0%)* |
| Diabetes mellitus | 827 (27.9%) | 14 (13.5%) | 36 (48.6%) | 7 (25.9%) | 11 (11.5%) | 13 (14.9%) | 52 (29.4%) | 66 (21.1%) | 195 (30.5%) | 18 (13.0%) | 12 (13.3%) | 246 (32.7%) | 151 (36.8%) | 6 (10.7%) |
| *Missing* | *70 (2.3%)* | *0* | *0* | *0* | *1 (1.0%)* | *0* | *0* | *0* | *2 (0.3%)* | *0* | *10 (10.0%)* | *0* | *0* | *57 (50.4%)* |
| **Cognitive outcomes** | | | | | | | | | | | | | | |
| Any-domain PSCI | 1,306 (43.0%) | 41 (39.4%) | 31 (41.9%) | 12 (44.4%) | 11 (11.3%) | 22 (25.3%) | 63 (35.6%) | 186 (59.4%) | 290 (45.2%) | 25 (18.1%) | 3 (3.0%) | 469 (62.3%) | 84 (20.5%) | 69 (61.1%) |
| Impaired attention / executive function | 377 (16.5%) | 9 (8.7%) | 4 (5.4%) | 2 (7.4%) | — | 2 (2.3%) | 16 (9.0%) | 119 (38.1%) | 50 (8.4%) | 7 (5.1%) | — | 149 (22.8%) | — | 19 (17.1%) |
| *Missing* | *755 (24.9%)* | *0* | *0* | *0* | *97 (100.0%)* | *0* | *0* | *1 (0.3%)* | *44 (6.9%)* | *1 (0.7%)* | *100 (100.0%)* | *100 (13.3%)* | *410 (100.0%)* | *2 (1.8%)* |
| Impaired processing speed | 362 (16.6%) | 27 (26.0%) | 22 (29.7%) | 9 (33.3%) | — | 2 (2.3%) | 10 (5.6%) | 106 (34.2%) | 44 (7.3%) | 8 (5.9%) | — | 134 (20.2%) | — | — |

| Characteristic | Pooled | CASPER | COAST | CODECS | CROMIS-2 | DEDEMAS | PROCRAS | GRECogVASC | Hallym VCI | STROKDEM | MSS-2 | Bundang VCI | CU-STRIDE | USCOG |
|---|---|---|---|---|---|---|---|---|---|---|---|---|---|---|
| *Missing* | *859 (28.3%)* | *0* | *0* | *0* | *97 (100.0%)* | *0* | *0* | *3 (1.0%)* | *42 (6.6%)* | *3 (2.2%)* | *100 (100.0%)* | *91 (12.1%)* | *410 (100.0%)* | *113 (100.0%)* |
| Impaired language | 373 (15.6%) | 6 (7.7%) | 2 (2.7%) | 5 (19.2%) | — | 1 (1.1%) | 19 (10.9%) | 119 (38.0%) | 39 (6.1%) | 2 (1.5%) | — | 160 (21.4%) | — | 20 (17.7%) |
| *Missing* | *646 (21.3%)* | *26 (25.0%)* | *0* | *1 (3.7%)* | *97 (100.0%)* | *0* | *2 (1.1%)* | *0* | *3 (0.5%)* | *1 (0.7%)* | *100 (100.0%)* | *6 (0.8%)* | *410 (100.0%)* | *0* |
| Impaired verbal memory | 611 (25.8%) | 9 (8.7%) | 7 (9.5%) | — | — | 6 (6.9%) | 46 (26.0%) | 48 (15.3%) | 130 (20.4%) | 12 (8.8%) | — | 313 (42.2%) | — | 40 (39.6%) |
| *Missing* | *664 (21.9%)* | *0* | *0* | *27 (100.0%)* | *97 (100.0%)* | *0* | *0* | *0* | *4 (0.6%)* | *2 (1.4%)* | *100 (100.0%)* | *12 (1.6%)* | *410 (100.0%)* | *12 (10.6%)* |
| Impaired visuospatial function | 522 (26.7%) | — | 16 (21.6%) | — | — | 13 (15.1%) | — | 57 (18.8%) | 135 (23.0%) | 7 (5.2%) | — | 269 (40.4%) | — | 25 (23.1%) |
| *Missing* | *1,076 (35.5%)* | *104 (100.0%)* | *0* | *27 (100.0%)* | *97 (100.0%)* | *1 (1.1%)* | *177 (100.0%)* | *10 (3.2%)* | *54 (8.4%)* | *4 (2.9%)* | *100 (100.0%)* | *87 (11.6%)* | *410 (100.0%)* | *5 (4.4%)* |
| Impaired visuospatial memory | 98 (11.0%) | — | 7 (9.5%) | — | — | 0 (0.0%) | — | — | 89 (14.7%) | 2 (1.6%) | — | — | — | — |

Continuous variables are given as median [interquartile range] and mean ± standard deviation. Categorical variables are given as n (%), with percentages calculated among patients with available data for that variable.

Missing indicates the number (%) of patients without available data for the preceding variable; 0 denotes complete data. An em dash (—) indicates that a variable or category was not assessed or not present in that cohort.

NPA = neuropsychological assessment; NIHSS = National Institutes of Health Stroke Scale; PSCI = post-stroke cognitive impairment; TIA = transient ischaemic attack

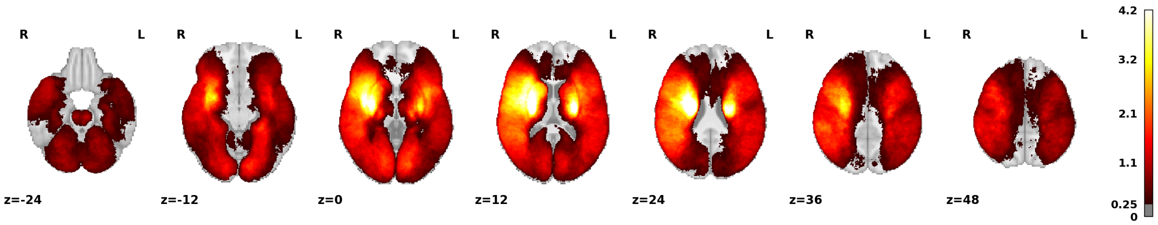


**Figure S1. Distribution of acute infarcts.** Voxelwise frequency of acute infarcts across the analysed sample (n = 3,034), shown on axial slices in MNI152NLin6Asym space in radiological convention. The colour scale indicates the percentage of patients with an infarct in each voxel.

L = left; R = right.

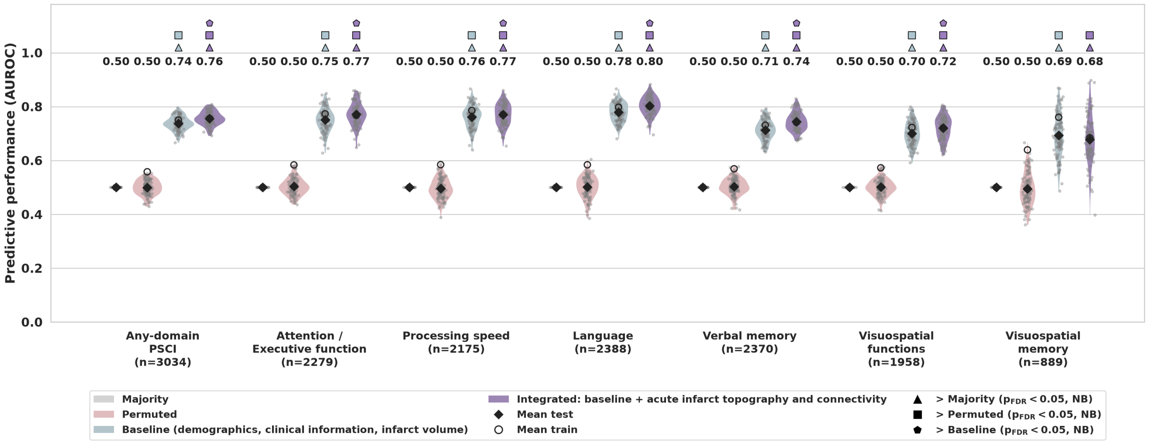


**Figure S2. Prediction of PSCI based on acute infarct topography and connectivity, demographics and clinical information — AUROC.** Predictive performance for each cognitive target based on pooled 10 × 10 repeated stratified cross-validation, quantified by the area under the receiver operating characteristic curve (AUROC) as a complement to the AUPRC reported in the main manuscript. Four models are compared: majority (prevalence-level reference), permuted (model trained on shuffled labels), baseline (demographics, clinical information, and infarct volume), and integrated (baseline plus acute infarct topography and connectivity: lesion location, functional LNM, and structural LNM scores). Diamonds indicate mean test AUROC across folds, with the value shown above each model"s violin; open circles indicate mean train AUROC. Symbols above a violin indicate that the model performed significantly better than a reference model (FDR-corrected Nadeau–Bengio paired t-tests, $p_{FDR}$ < 0.05): triangle, better than majority; square, better than permuted; pentagon, better than baseline. Consistent with the AUPRC results, the integrated model performed significantly better than the baseline for six of the seven cognitive targets, with visuospatial memory again the exception. Majority and permuted references remained at chance level (AUROC ≈ 0.50).

AUROC = area under the receiver operating characteristic curve; AUPRC = area under the precision–recall curve; PSCI = post-stroke cognitive impairment; functional LNM = functional lesion network mapping; structural LNM = structural lesion network mapping; NB = Nadeau–Bengio; FDR = false discovery rate.

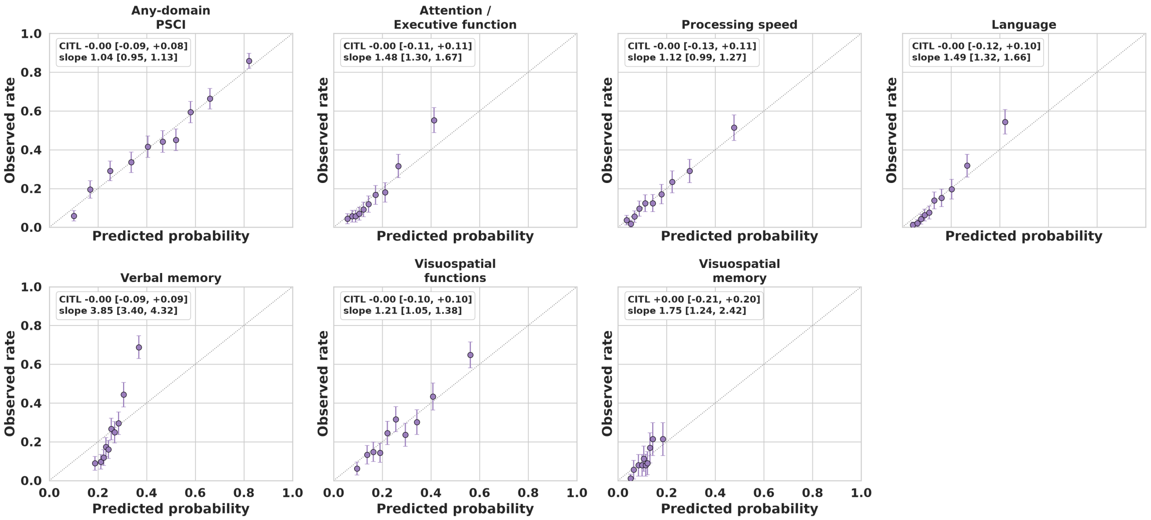


**Figure S3. Calibration of the integrated model.** Observed impairment rate plotted against the mean predicted probability of the integrated model within deciles of predicted risk, based on out-of-fold predictions from pooled 10 × 10 repeated stratified cross-validation, averaged per patient across the 10 repeats. Error bars denote the binomial standard error; the dotted diagonal indicates perfect calibration. Each panel reports calibration-in-the-large (CITL), estimated as the intercept of a logistic regression of the observed outcome with the logit of the predicted probability as offset (ideal value 0), and the calibration slope, estimated as the coefficient of a logistic regression of the observed outcome on the logit of the predicted probability (ideal value 1), each with 95% confidence intervals from 1,000 subject-level bootstrap resamples. Predicted risks were correct on average for every target (CITL between −0.005 and 0.004; all 95% confidence intervals included 0). The calibration slope was 1.04 (95% CI 0.95–1.13) for any-domain PSCI and ranged from 1.12 (0.99–1.27) for processing speed to 3.85 (3.40–4.32) for verbal memory; the confidence interval excluded 1 for all domains except processing speed. Slopes above 1 indicate that predicted probabilities were less dispersed than the observed risks, i.e. conservative rather than overconfident, consistent with shrinkage by the regularised base and meta-learners and by averaging across repeats. Predicted probabilities, particularly for verbal memory, would therefore require recalibration before use as absolute risk estimates, whereas the ranking of patients evaluated by AUPRC and AUROC is unaffected.

AUROC = area under the receiver operating characteristic curve; AUPRC = area under the precision–recall curve; CI = confidence interval; CITL = calibration-in-the-large; PSCI = post-stroke cognitive impairment.

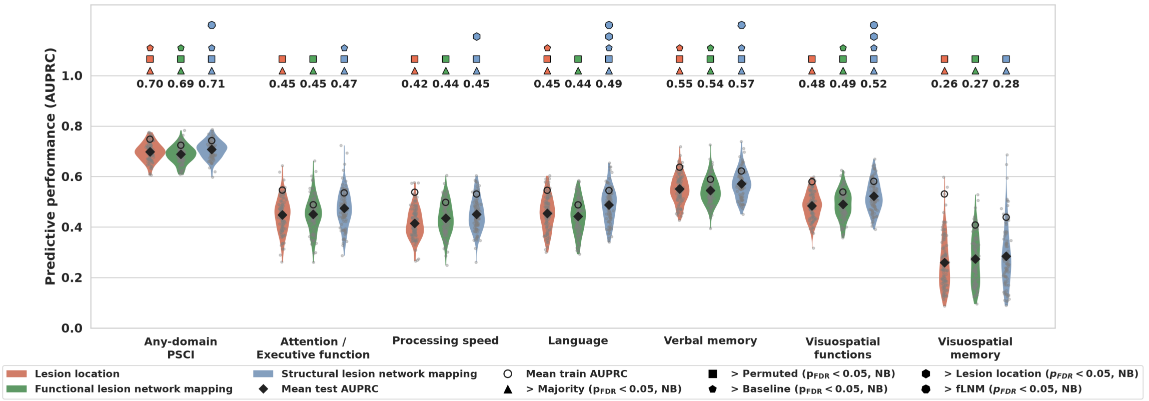


**Figure S4. Predictive performance of individual acute infarct topography and connectivity representations.** Predictive performance for each cognitive target based on pooled 10 × 10 repeated stratified cross-validation. Three single-representation models are compared, each combining the baseline variables (demographics, clinical information, and infarct volume) with one representation of the acute infarct: lesion location (red), functional LNM (green), or structural LNM (blue). Violins show the distribution of test AUPRC across the 100 outer folds; diamonds indicate mean test AUPRC across folds, with the value printed above each model; open circles indicate mean train AUPRC across folds. Symbols above a model's violin, in that model's colour, indicate that it performed significantly better than a reference model (FDR-corrected Nadeau–Bengio paired t-tests, $p_{FDR} < 0.05$): triangle, better than majority; square, better than permuted; pentagon, better than baseline; hexagon, better than lesion location; heptagon, better than functional LNM. The majority-class and permuted-label reference models are not shown but were evaluated on identical fold assignments. Structural LNM achieved the highest mean test AUPRC for every cognitive target and significantly outperformed lesion location (3 of 7 targets) and functional LNM (4 of 7 targets). This is consistent with the ablation analysis in the main manuscript, where performance of the integrated model dropped when structural LNM was removed but was largely unaffected by removal of lesion location or functional LNM.

AUPRC = area under the precision–recall curve; functional LNM = functional lesion network mapping; structural LNM = structural lesion network mapping; NB = Nadeau–Bengio; FDR = false discovery rate.

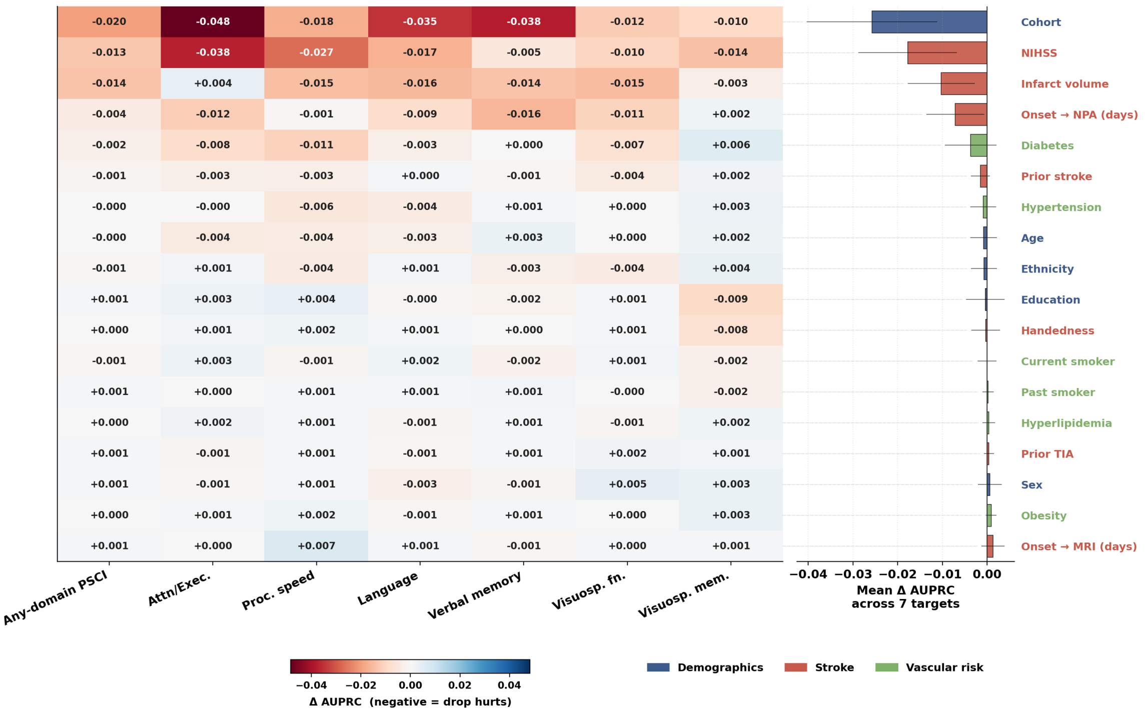


**Figure S5. Baseline variable ablation analysis across cognitive outcomes.** The figure illustrates the impact of removing individual baseline variables on the predictive performance of the baseline model across the seven cognitive targets. Performance changes are quantified as ΔAUPRC based on a pooled 10 × 10 repeated stratified cross-validation. Left (Heatmap): Displays the specific ΔAUPRC when each variable is dropped for a given cognitive domain. Negative values (red shading) indicate that removing the variable decreases predictive performance, signifying that the variable contributes information to the model. Right (Bar Plot): A horizontal bar chart showing the mean ΔAUPRC (with error bars indicating standard deviation) across all seven cognitive targets for each ablated variable. Variables are ranked top-to-bottom by their mean ΔAUPRC and are colour-coded by category: demographics (blue), clinical stroke information (red), and vascular risk factors (green). Removing cohort membership, NIHSS, Infarct volume and the time between stroke onset and neuropsychological assessment from the baseline model led to strongest reductions in predictive performance.

Attn/Exec. = Attention / executive function. AUPRC = area under the precision–recall curve; NIHSS = National Institutes of Health Stroke Scale; NPA = neuropsychological assessment; MRI = magnetic resonance imaging; proc. speed = processing speed; PSCI = post-stroke cognitive impairment; TIA = transient ischemic attack; visuosp. fn. = visuospatial function; visuosp. mem. = visuospatial memory

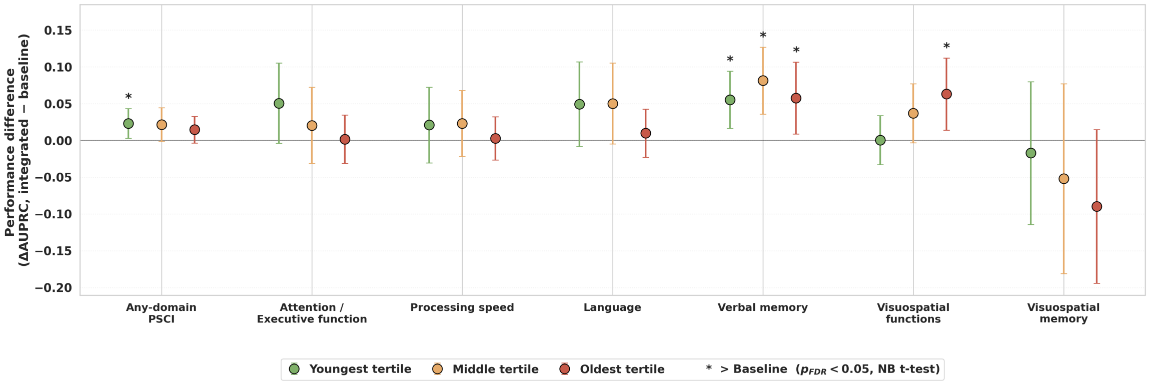


**Figure S6. Benefit of the integrated model stratified by age.** Performance difference (ΔAUPRC, integrated minus baseline) for each cognitive target within age tertiles (youngest: < 63 years, n = 999; middle: 63 to < 73 years, n = 954; oldest: ≥ 73 years, n = 1,081). Models were refitted within each tertile under pooled 10 × 10 repeated stratified cross-validation, with age removed from the baseline variables. Points show the mean fold-paired difference across the 100 outer folds; whiskers denote the 95% confidence intervals based on the Nadeau-Bengio corrected variance. Asterisks denote integrated > baseline (one-sided Nadeau–Bengio t-tests, FDR-corrected across the three age tertiles within each cognitive target, $p_{\mathrm{FDR}} < 0.05$).

AUPRC = area under the precision–recall curve; FDR = false discovery rate; NB = Nadeau–Bengio; PSCI = post-stroke cognitive impairment; SE = standard error.

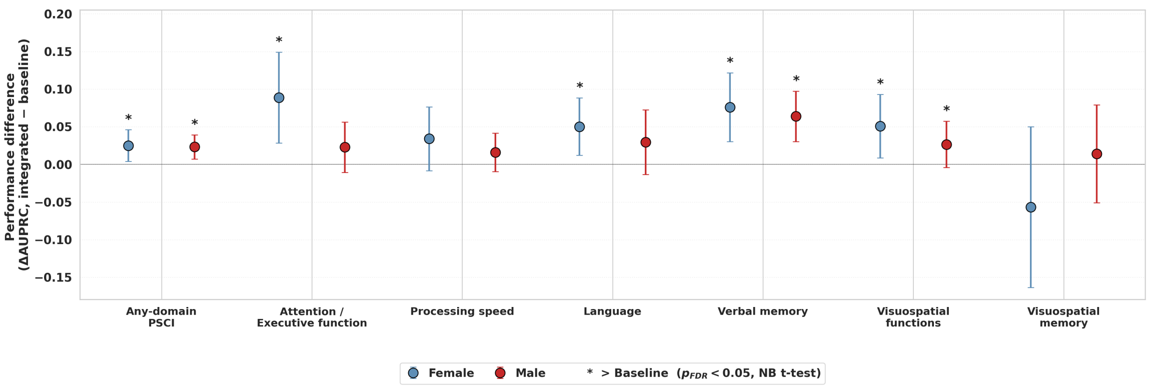


**Figure S7. Benefit of the integrated model stratified by sex.** Performance difference (ΔAUPRC, integrated minus baseline) for each cognitive target within women (n = 1,184) and men (n = 1,850). Models were refitted within each stratum under pooled 10 × 10 repeated stratified cross-validation, with sex removed from the input features. Points show the mean fold-paired difference across the 100 outer folds; whiskers denote the 95% confidence intervals based on the Nadeau-Bengio corrected variance. Asterisks denote integrated > baseline (one-sided Nadeau–Bengio t-tests, FDR-corrected across the two sex strata, $p_{\mathrm{FDR}} < 0.05$).

AUPRC = area under the precision–recall curve; FDR = false discovery rate; NB = Nadeau–Bengio; PSCI = post-stroke cognitive impairment; SE = standard error.

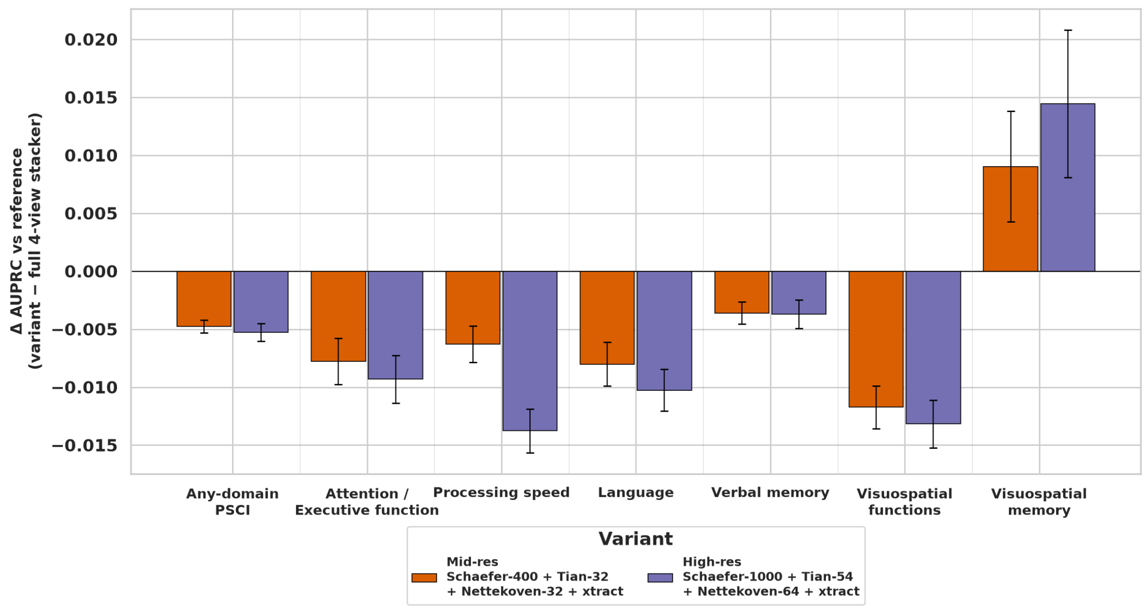


**Figure S8. Sensitivity of predictive performance to parcellation resolution.** ΔAUPRC (variant minus reference) of the integrated model for each cognitive target, comparing the 190-region reference parcellation (Schaefer-100 cortex, Tian S1 subcortex, Nettekoven-32 cerebellum, XTRACT white matter) with a mid-resolution (Schaefer-400, Tian S2, Nettekoven-32, XTRACT) and a high-resolution variant (Schaefer-1000, Tian S4, Nettekoven-64, XTRACT). All models were evaluated on identical folds under pooled 10 × 10 repeated stratified cross-validation. Bars show the mean difference across the 100 outer folds; error bars denote the across-fold SE. Finer parcellations reduced predictive performance for six of seven targets, with an improvement only for visuospatial memory.

AUPRC = area under the precision–recall curve; PSCI = post-stroke cognitive impairment; SE = standard error.

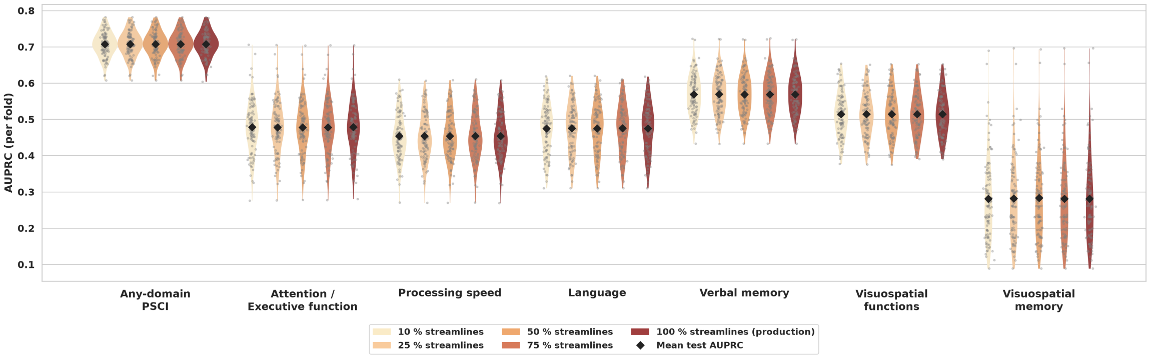


**Figure S9. Sensitivity of predictive performance to normative tractogram density.** Test AUPRC of the integrated model for each cognitive target, with structural LNM scores derived from random subsamples of 10%, 25%, 50%, 75% and 100% of the dTOR985 streamlines (100% used in the main analysis). All variants were evaluated on identical folds under pooled 10 × 10 repeated stratified cross-validation. Violins show the distribution of test AUPRC across the 100 outer folds; diamonds indicate the mean. Mean performance was unchanged across subsampling fractions for every target.

AUPRC = area under the precision–recall curve; dTOR985 = normative structural connectome of 985 subjects; LNM = lesion network mapping; PSCI = post-stroke cognitive impairment.

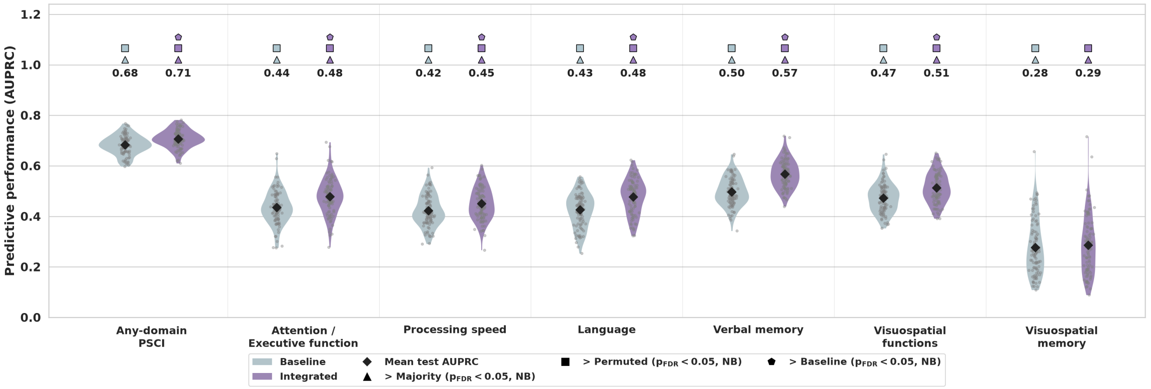


**Figure S10. Predictive performance with log-transformed infarct volume.** Test AUPRC of the baseline and integrated models under pooled 10 × 10 repeated stratified cross-validation with log(1 + infarct volume) replacing raw infarct volume in all models. Violins and grey dots show the distribution of test AUPRC across the 100 outer folds; diamonds indicate the mean, printed above each violin. Symbols above the violins indicate that the model performed significantly better than a reference model (FDR-corrected one-sided Nadeau–Bengio paired t-tests, $p_{FDR}$ < 0.05): triangle, better than majority; square, better than permuted; pentagon, better than baseline. Majority and permuted models were evaluated on identical folds but are not shown. As in the main analysis, the integrated model improved on the baseline for six of seven targets. Compared with the main analysis on identical folds, log transformation changed mean AUPRC by at most 0.004 for the integrated model and at most 0.009 for the baseline.

AUPRC = area under the precision–recall curve; FDR = false discovery rate; NB = Nadeau–Bengio; PSCI = post-stroke cognitive impairment.

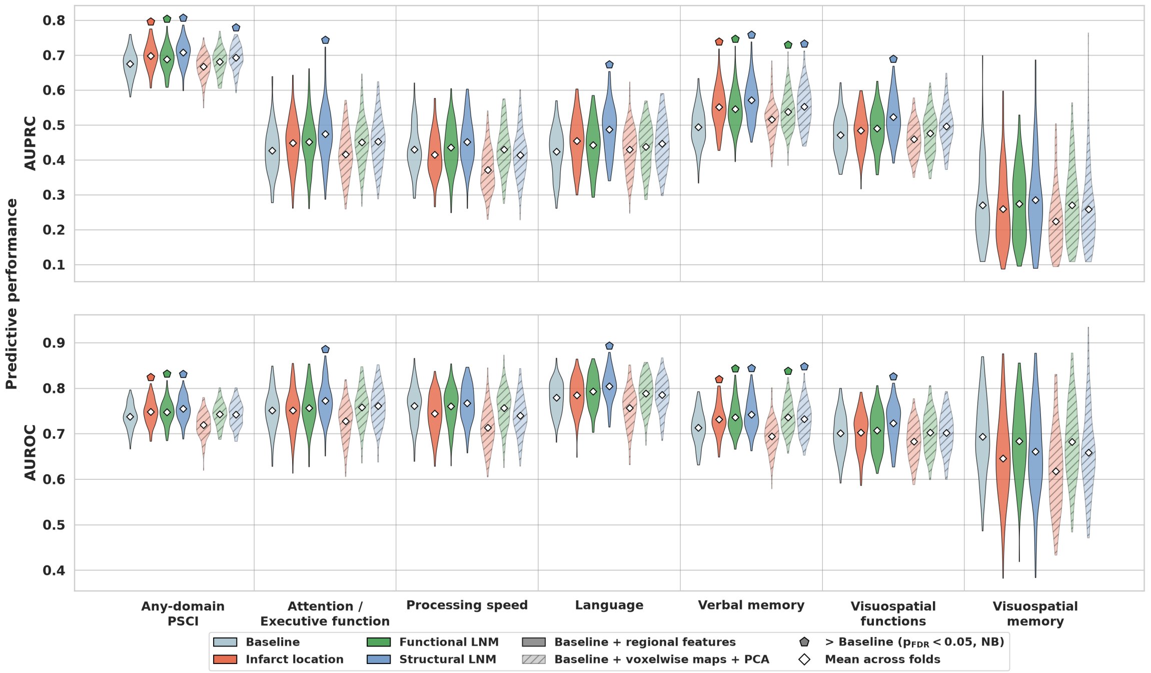


**Figure S11. Regional versus voxelwise lesion features.** Out-of-sample AUPRC (top) and AUROC (bottom) of single-representation models (baseline plus one acute infarct representation) under pooled 10 × 10 repeated stratified cross-validation, with lesion location (red), functional LNM (green) or structural LNM (blue) entered either as scores in the 190 regions of the reference parcellation (left, solid) or as voxelwise maps at 2 mm resolution reduced to the principal components explaining 95% of their variance (right, hatched). The principal component analysis was fitted within each training fold and applied to the corresponding test fold. Violins show the distribution across the 100 outer folds and white diamonds the mean. Pentagons mark variants that outperformed the baseline model (one-sided Nadeau–Bengio corrected t-tests, FDR-corrected across the six variants within each target and metric; pFDR < 0.05).

AUPRC = area under the precision–recall curve; AUROC = area under the receiver operating characteristic curve; LNM = lesion network mapping; PSCI = post-stroke cognitive impairment.

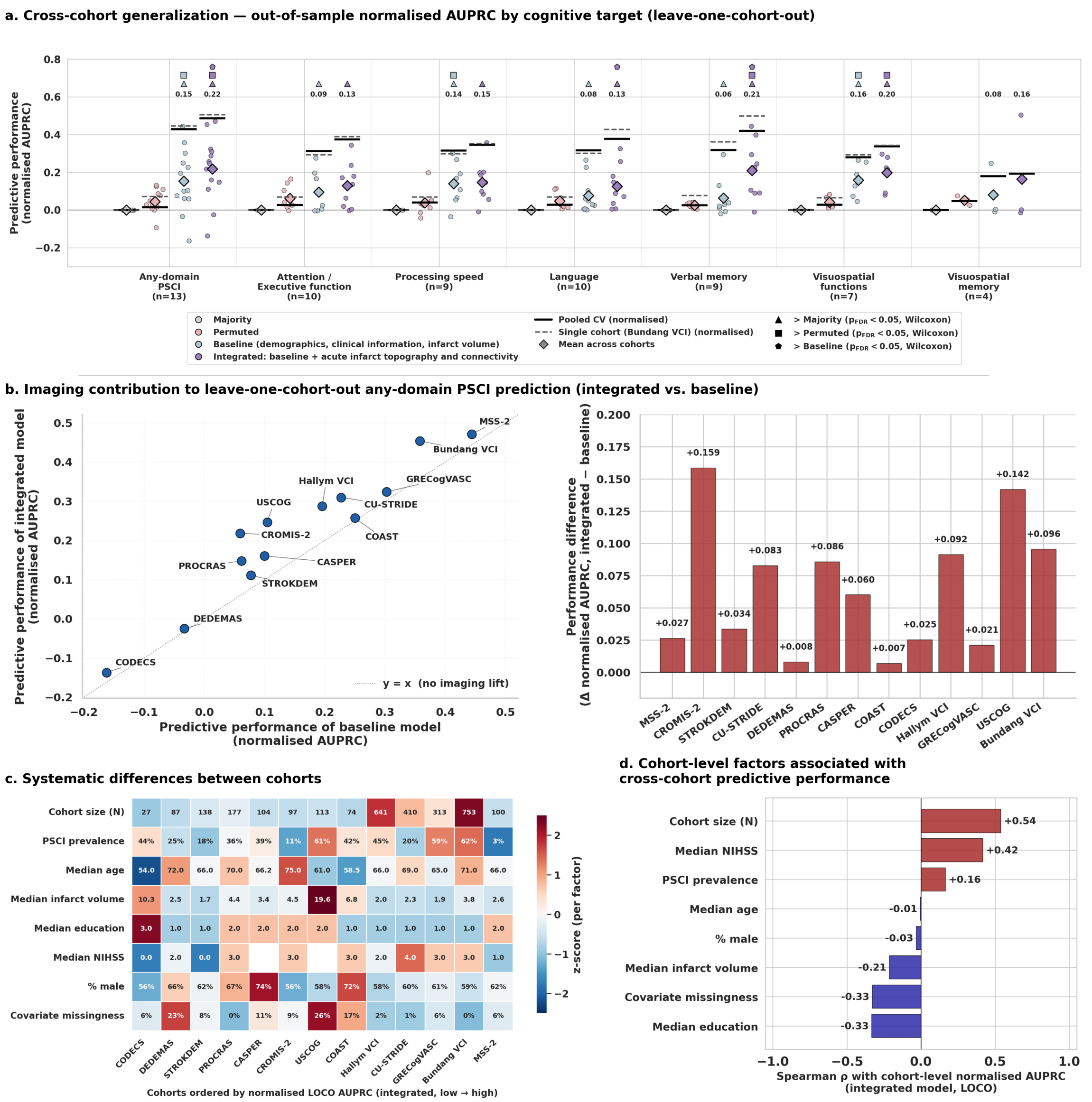


**Figure S12. Cross-cohort generalisation under leave-one-cohort-out cross-validation. (a)** Predictive performance for each cognitive target based on leave-one-cohort-out cross-validation to assess cross-cohort generalisation. Performance is measured based on out-of-sample normalised AUPRC defined as (AUPRC − prevalence) / (1 − prevalence) to account for between-cohort prevalence differences (chance level = 0). Four models were compared: majority (prevalence-level baseline), permuted (model trained on shuffled labels), “baseline” (demographics, clinical information, and infarct volume), and integrated (baseline plus multi-modal acute infarct topography: lesion location, functional LNM, and structural LNM). Each dot represents a single held-out cohort; diamonds indicate the mean across cohorts. Horizontal lines show the pooled 10x10 CV reported in the main manuscript (solid) and single-cohort (Bundang VCI; dashed) reference performance. Symbols above a violin indicate that the model performed significantly better than a reference model (FDR-corrected Wilcoxon signed-rank tests across held-out cohorts, $p_{FDR} < 0.05$): triangle, better than majority; square, better than permuted; pentagon, better than baseline. The number of cohorts contributing data per domain is shown in parentheses. The integrated model exhibited significantly higher predictive performance than the baseline for three of seven cognitive targets. **(b)** Contribution of infarct topography and connectivity to any-domain PSCI prediction in each held-out cohort. **Left:** normalised AUPRC of the integrated model plotted against the baseline model, with one point per cohort. Points above the diagonal (y = x) indicate cohorts in which adding infarct topography and connectivity improved prediction over demographics, clinical information, and infarct volume alone. **Right:** the same information expressed as the per-cohort difference in normalised AUPRC (integrated − baseline). All 13 cohorts showed a positive contribution, indicating that the predictive benefit was not driven by individual cohorts. **(c)** Systematic differences between cohorts. Cohort-level characteristics (cohort size, any-domain PSCI prevalence, median age, median infarct volume, median education, median NIHSS, proportion of male patients and proportion of missing baseline covariate values), coloured by per-characteristic z-score across cohorts and annotated with raw values. Cohorts are ordered by the normalised AUPRC of the integrated model for any-domain PSCI; NIHSS was not available in two cohorts. **(d)** Spearman correlation of each characteristic with the cohort-level normalised AUPRC of the integrated model for any-domain PSCI.

AUPRC = area under the precision–recall curve; FDR = false discovery rate; NIHSS = National Institutes of Health Stroke Scale; LOCO = leave-one-cohort-out; PSCI = post-stroke cognitive impairment.

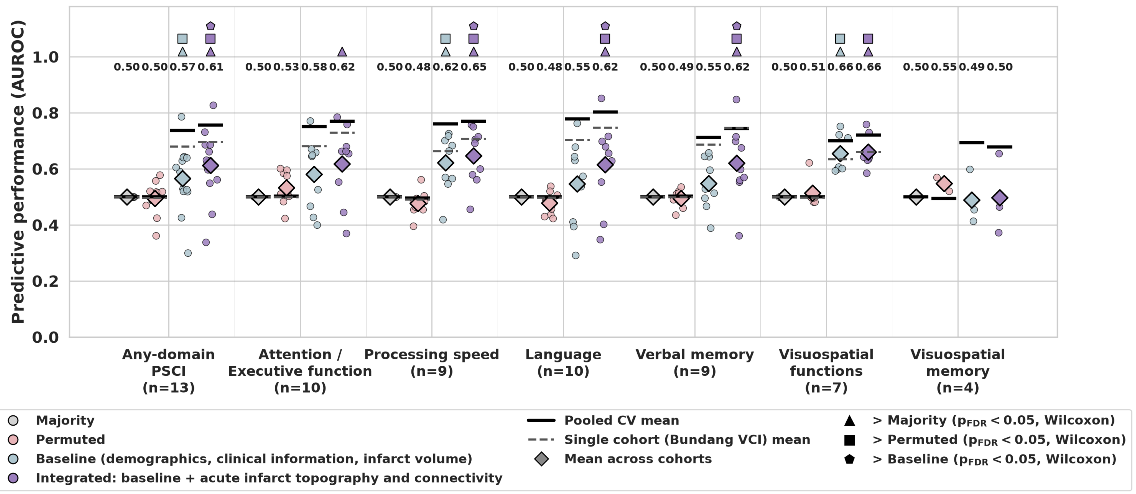


**Figure S13. Cross-cohort generalisation of PSCI prediction — out-of-sample AUROC by cognitive target (leave-one-cohort-out).** Predictive performance under leave-one-cohort-out cross-validation, in which each cohort served once as an external test set, quantified by the area under the receiver operating characteristic curve (AUROC). Because AUROC has a fixed chance level of 0.50 irrespective of impairment prevalence, no prevalence normalisation was applied. Four models are compared: majority (chance-level reference), permuted (model trained on shuffled labels), baseline (demographics, clinical information, and infarct volume), and integrated (baseline plus acute infarct topography and connectivity). Small circles indicate individual held-out cohorts; diamonds indicate mean test AUROC across cohorts; solid horizontal lines indicate the pooled cross-validation mean; dashed lines indicate the largest single-cohort (Bundang VCI) mean. Numbers above each target denote mean AUROC for the baseline and integrated models. Symbols above the violins indicate that the model performed significantly better than a reference model (FDR-corrected paired Wilcoxon signed-rank tests across held-out cohorts, $p_{FDR} < 0.05$): triangle, better than majority; square, better than permuted; pentagon, better than baseline.The integrated model was significantly better than the clinical baseline for any-domain PSCI, processing speed, language and verbal memory. n denotes the number of cohorts contributing to each cognitive target.

AUROC = area under the receiver operating characteristic curve; AUPRC = area under the precision–recall curve; PSCI = post-stroke cognitive impairment; CV = cross-validation; FDR = false discovery rate.

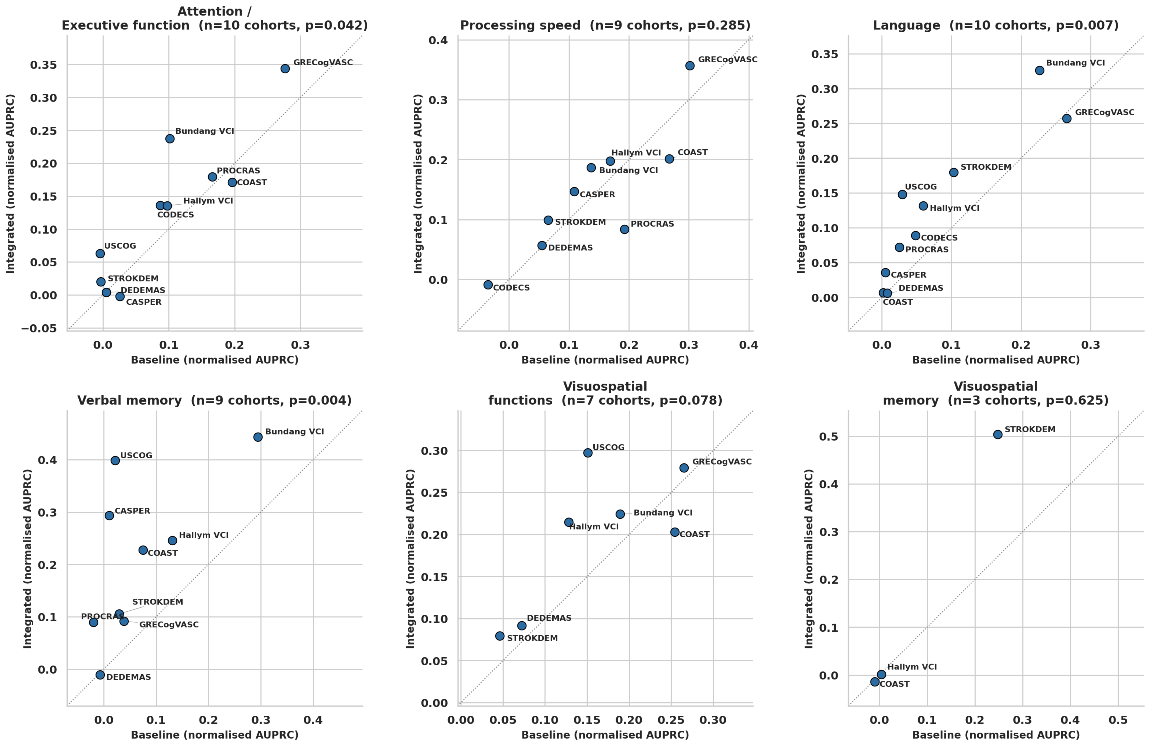


**Figure S14. Cross-cohort transfer for individual cognitive domains.** For each cognitive domain, normalised AUPRC of the integrated model is plotted against the baseline model under leave-one-cohort-out cross-validation, with one point per held-out cohort contributing data for that domain. Points above the diagonal (y = x) indicate cohorts in which acute infarct topography and connectivity improved prediction. Panel titles give the number of cohorts and the uncorrected P value of a one-sided paired Wilcoxon signed-rank test (integrated > baseline). After FDR correction across the six domains, the benefit was significant for language and verbal memory ($p_{FDR} < 0.05$ each).

AUPRC = area under the precision–recall curve; FDR = false discovery rate; PSCI = post-stroke cognitive impairment.

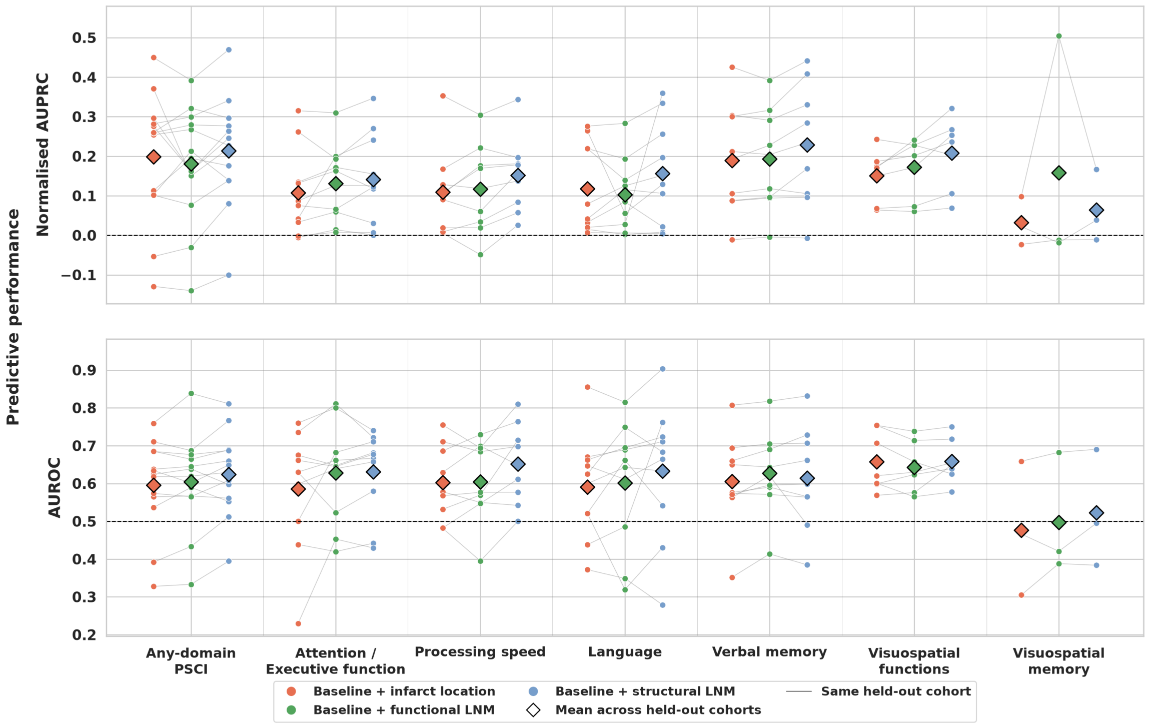


**Figure S15. Cross-cohort performance of the three lesion representations.** Out-of-sample performance under leave-one-cohort-out cross-validation of the baseline model extended by a single acute infarct representation: infarct location (red), functional LNM (green) or structural LNM (blue). Each point is one held-out cohort; grey lines connect the same held-out cohort across representations, and diamonds show the mean across held-out cohorts. The number of held-out cohorts varies by outcome according to the availability of domain-specific assessments. Top: Normalised AUPRC, (AUPRC − prevalence) / (1 − prevalence), with the prevalence of each held-out cohort taken from the majority-class model (0 corresponds to chance). Bottom: AUROC; the dashed line marks chance (0.5).

AUPRC = area under the precision–recall curve; AUROC = area under the receiver operating characteristic curve; FDR = false discovery rate; LNM = lesion network mapping; PSCI = post-stroke cognitive impairment.

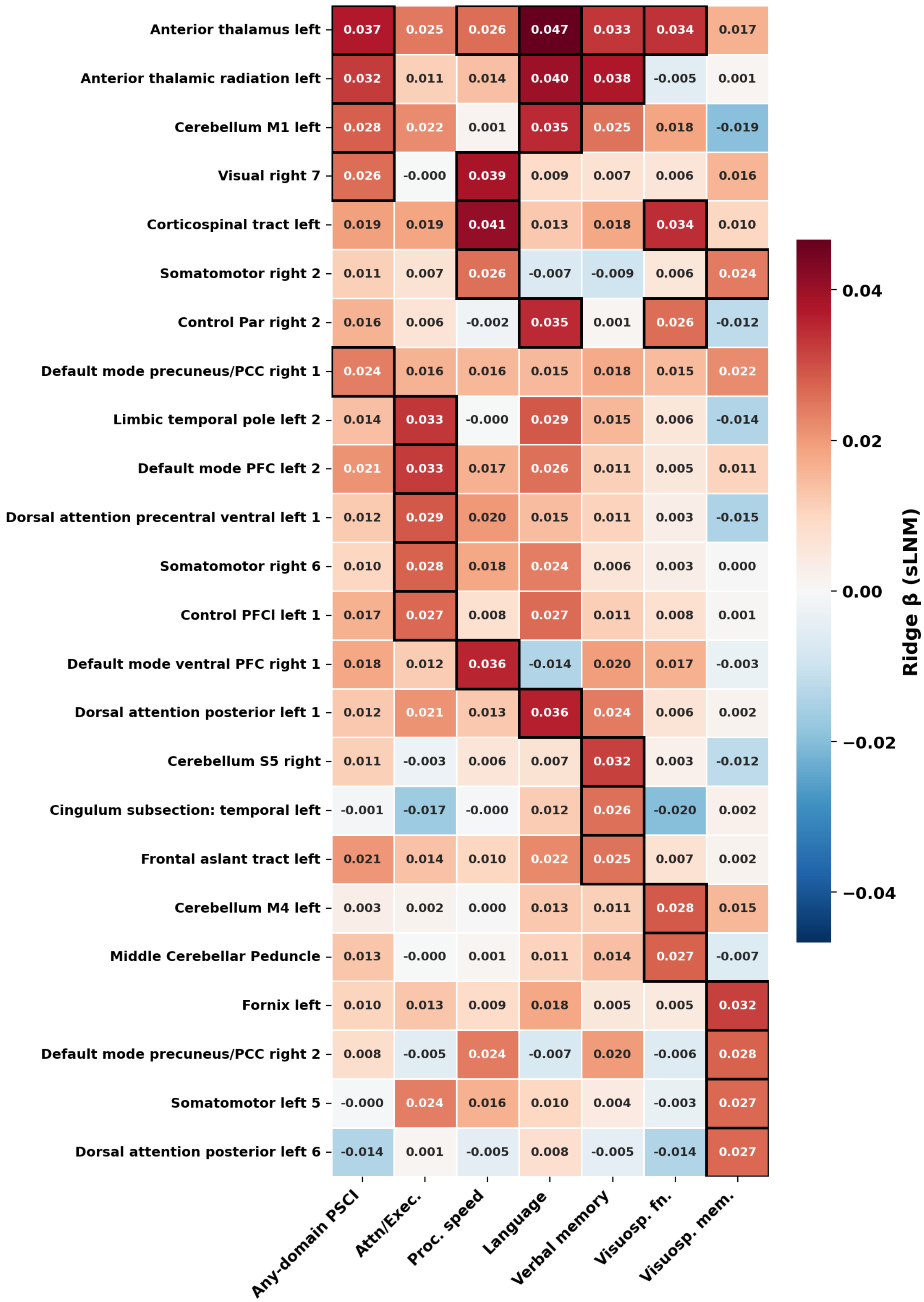


**Figure S16. Overlap of the most strongly weighted structural LNM regions across cognitive targets.** Rows comprise the union of the five regions with the largest stable positive coefficients (|mean β| / SE ≥ 2) of the structural LNM base learner for each cognitive target (24 regions). Cells show the mean ridge coefficient across the 100 pooled outer folds; black outlines mark regions among a target's top five. Rows are ordered by the number of targets for which a region ranks among the top five. Structural lesion connectivity of the left anterior thalamus ranked among the top five regions for five of seven targets.

Attn/Exec. = attention/executive function; LNM = lesion network mapping; PSCI = post-stroke cognitive impairment; SE = standard error.

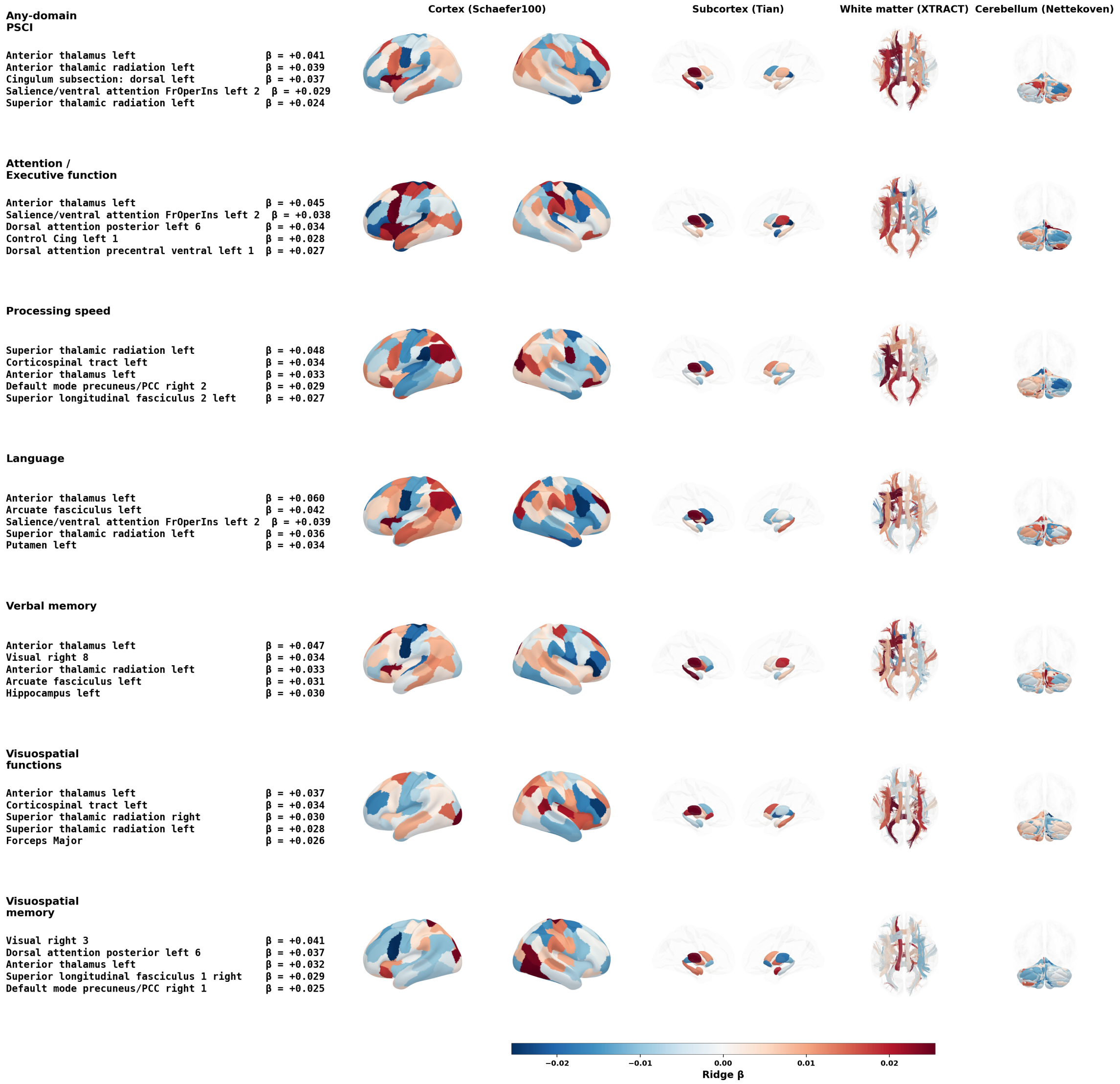


**Figure S17. Anatomy of cognitive domain-specific lesion location profiles.** As figure 3 of the main manuscript, for the base learner informed by lesion location scores. Standardised β coefficients were averaged across the 100 pooled outer cross-validation folds and displayed across cortical (Schaefer), subcortical (Tian), white matter (XTRACT) and cerebellar (Nettekoven) regions. Positive values (red) indicate that the model weights greater lesion overlap with a region towards a higher predicted probability of impairment. For each target, the five regions with the largest positive β values are listed on the left.

SE = standard error.

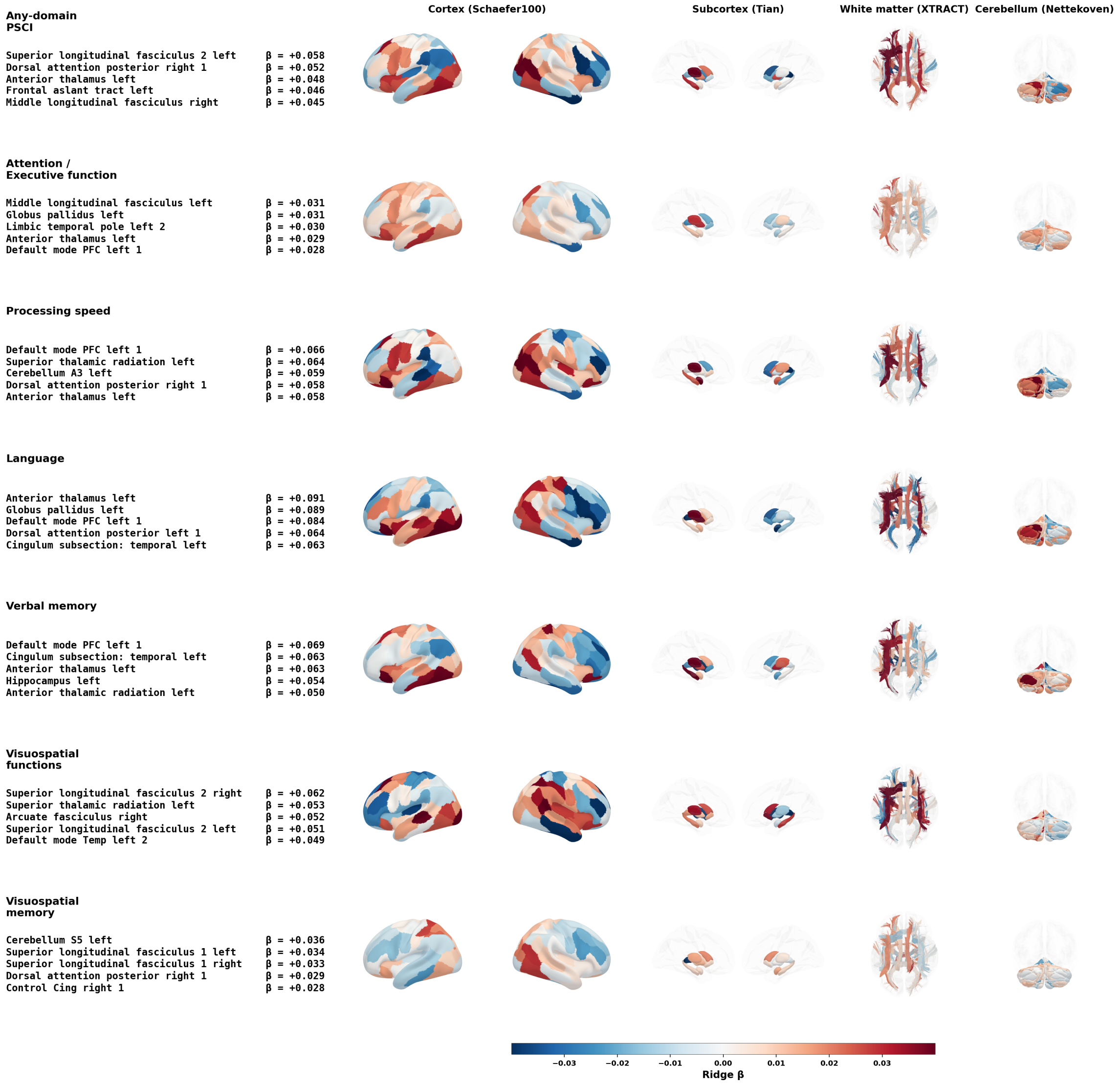


**Figure S18. Anatomy of cognitive domain-specific functional lesion network profiles.** As figure 3 of the main manuscript, for the base learner informed by functional LNM scores. Standardised β coefficients were averaged across the 100 pooled outer cross-validation folds and displayed across cortical (Schaefer), subcortical (Tian), white matter (XTRACT) and cerebellar (Nettekoven) regions. Positive values (red) indicate that the model weights greater functional connectivity of a region with the lesion towards a higher predicted probability of impairment. For each target, the five regions with the largest positive β values are listed on the left.

LNM = lesion network mapping; SE = standard error.

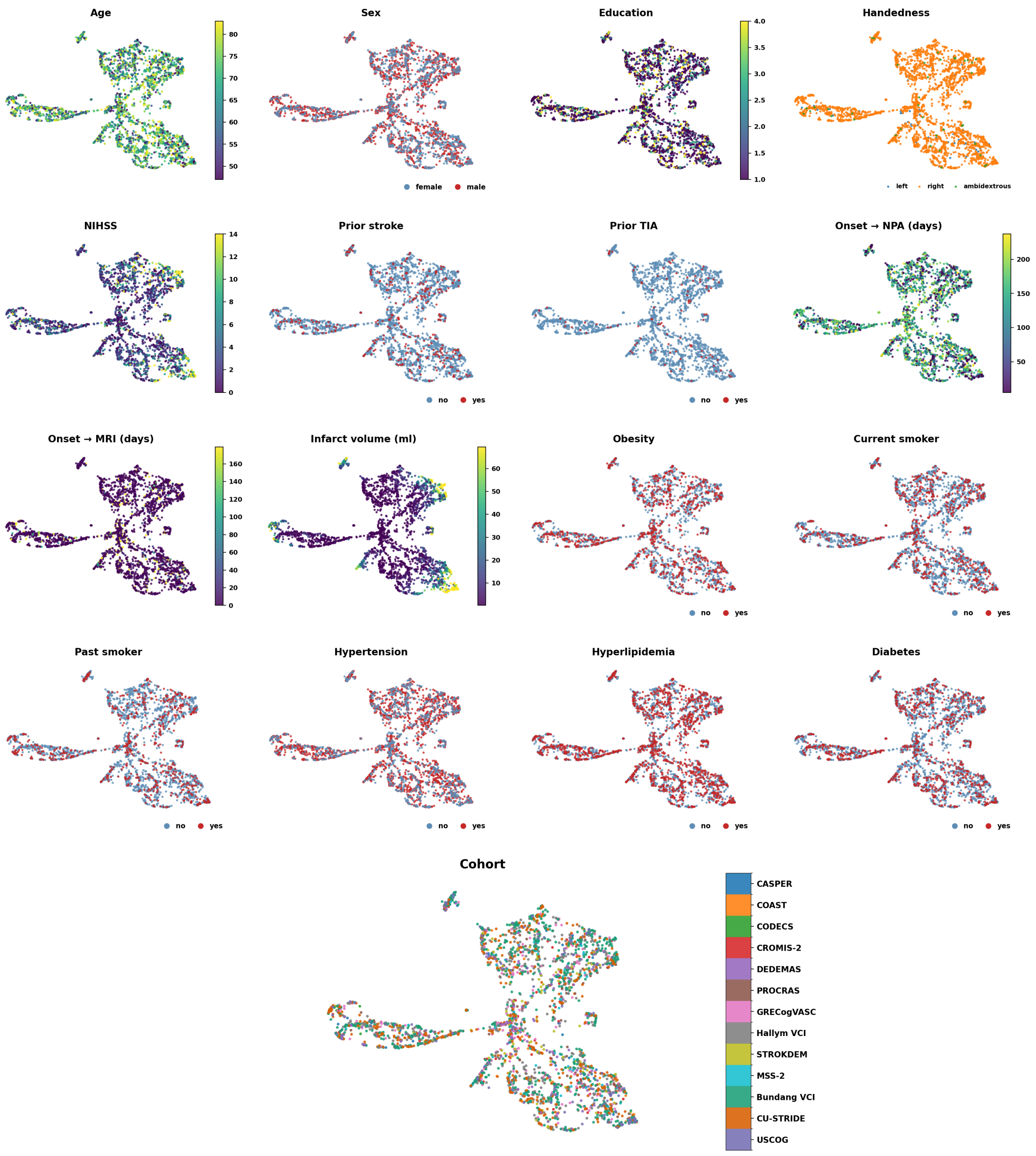


**Figure S19. Baseline covariates projected onto the structural LNM UMAP embedding.** Each panel shows the same UMAP embedding of parcel-level structural LNM scores, with patients coloured by a single demographic, clinical, or cohort characteristic. Infarct volume and NIHSS showed systematic structure across the embedding, with the highest values concentrated in the large left- and right-MCA clusters, whereas demographic characteristics, vascular risk factors, and cohort membership were distributed diffusely.

NPA = neuropsychological assessment; NIHSS = National Institutes of Health Stroke Scale; structural LNM = structural lesion network mapping; TIA = transient ischaemic attack; UMAP = uniform manifold approximation and projection.

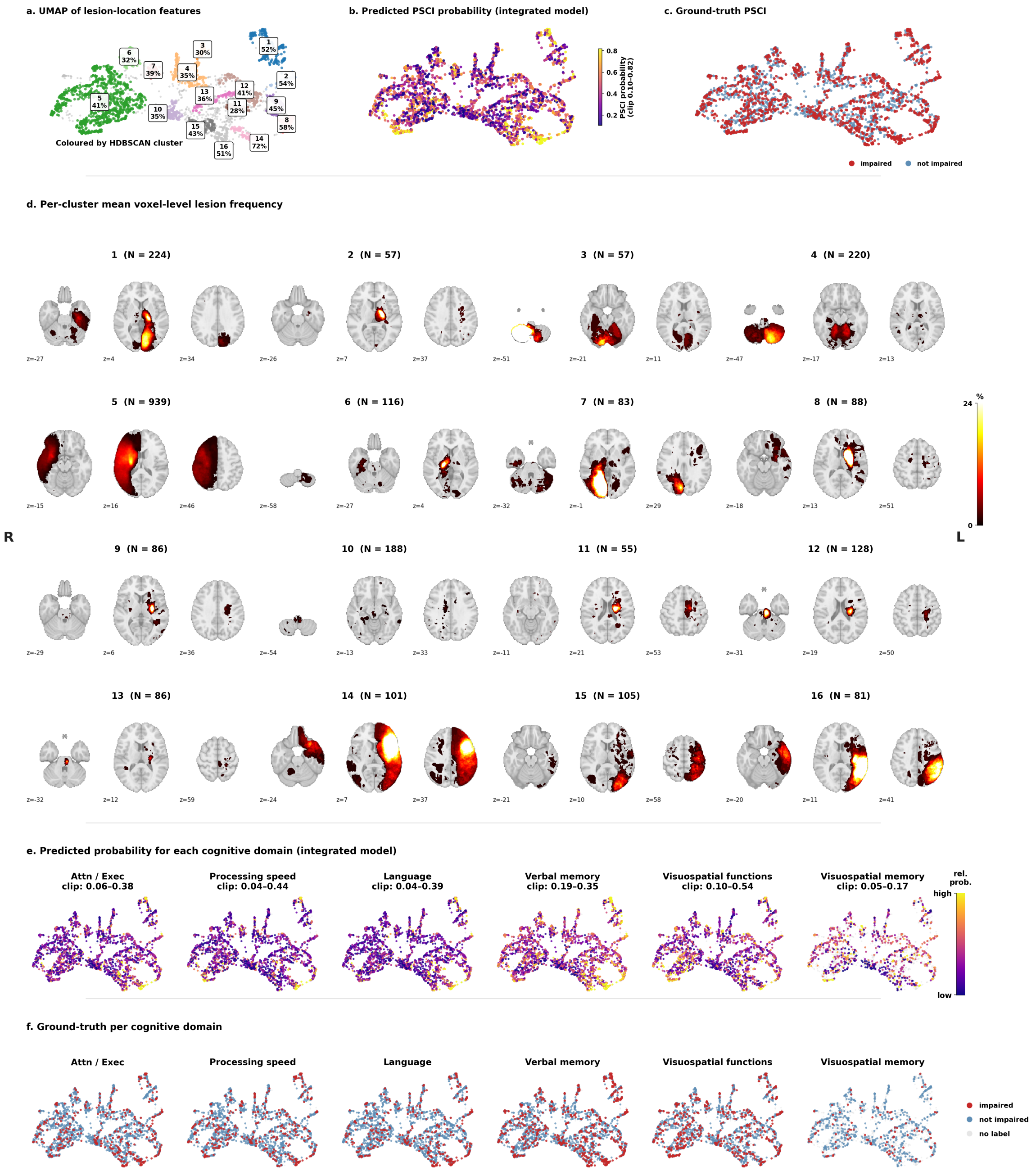


**Figure S20. Data-driven patient stratification by lesion location. (a)** UMAP embedding of parcel-level lesion location scores of acute infarcts, coloured by HDBSCAN cluster assignment. Sixteen clusters emerged, each labelled by observed any-domain PSCI prevalence. Of note, HDBSCAN does not necessarily assign every subject to a cluster; subjects not assigned to any cluster were labelled as noise and are displayed with reduced opacity. **(b)** Same UMAP embedding coloured by the integrated model's predicted any-domain PSCI probability. **(c)** Same embedding coloured by ground-truth any-domain PSCI status. **(d)** Per-cluster lesion frequency maps displayed at representative axial slices. The colour scale represents the percentage of patients exhibiting a lesion within each voxel. **(e)** Per-domain predicted impairment probability from the integrated model projected onto the UMAP embedding, with domain-specific colour scales clipped to the observed probability range. **(f)** Ground-truth impairment labels per cognitive domain projected onto the same embedding. Faded points indicate patients without available assessment for that domain.

Attn/Exec. = attention/executive function; HDBSCAN = hierarchical density-based spatial clustering of applications with noise; PSCI = post-stroke cognitive impairment; UMAP = uniform manifold approximation and projection.

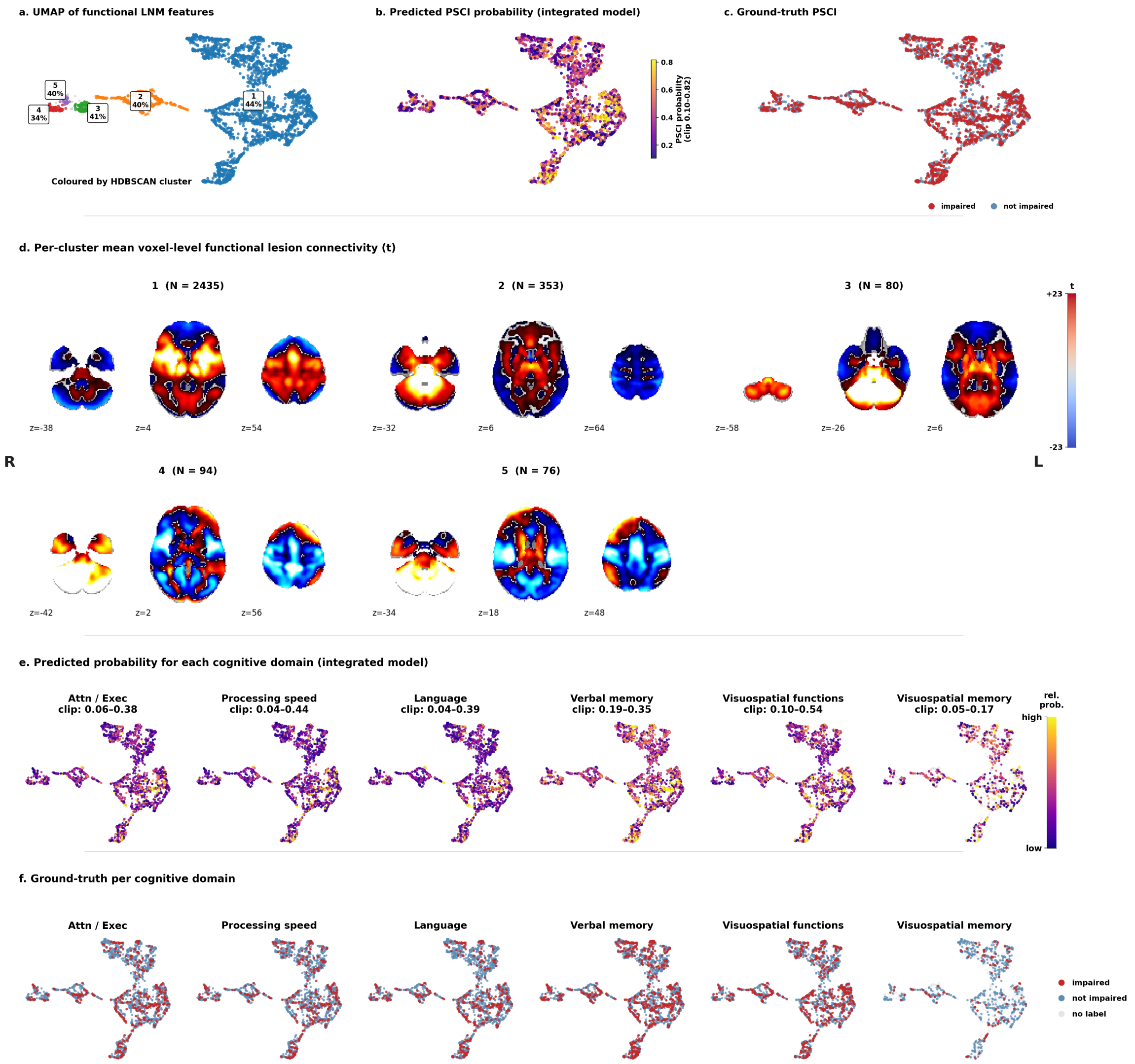


**Figure S21. Data-driven patient stratification by functional lesion network mapping.** **(a)** UMAP embedding of parcel-level functional lesion network mapping scores of acute infarcts, coloured by HDBSCAN cluster assignment. Five clusters emerged, each labelled by observed any-domain PSCI prevalence. Of note, HDBSCAN does not necessarily assign every subject to a cluster; subjects not assigned to any cluster were labelled as noise and are displayed with reduced opacity. **(b)** Same UMAP embedding coloured by the integrated model's predicted any-domain PSCI probability. **(c)** Same embedding coloured by ground-truth any-domain PSCI status. **(d)** Per-cluster lesion frequency maps displayed at representative axial slices, illustrating the anatomical distribution of areas functionally connected to the lesion. The colour scale represents the percentage of patients exhibiting a lesion within each voxel. **(e)** Per-domain predicted impairment probability from the integrated model projected onto the UMAP embedding, with domain-specific colour scales clipped to the observed probability range. **(f)** Ground-truth impairment labels per cognitive domain projected onto the same embedding. Faded points indicate patients without available assessment for that domain.

Attn/Exec. = attention/executive function; HDBSCAN = hierarchical density-based spatial clustering of applications with noise; LNM = lesion network mapping; PSCI = post-stroke cognitive impairment; UMAP = uniform manifold approximation and projection.

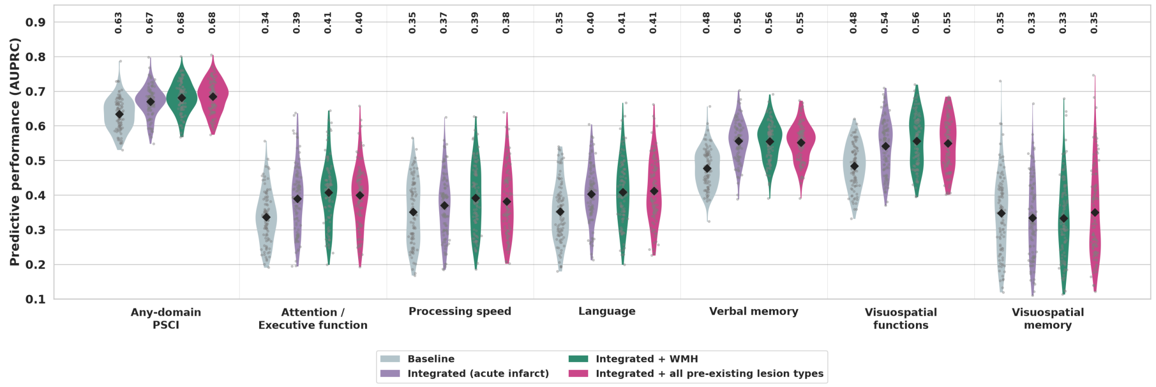


**Figure S22. Predictive performance with acute and pre-existing cerebrovascular lesion features.** Out-of-sample AUPRC of four models, evaluated by pooled 10 × 10 repeated stratified cross-validation in patients with segmentations of pre-existing cerebrovascular lesions: the baseline model (demographic and clinical variables and infarct volume); the integrated model (baseline plus acute infarct topography and connectivity); the integrated model plus WMH topography and connectivity; and the integrated model plus the topography and connectivity of all pre-existing lesion types combined (WMH, old infarcts and lacunes). Each pre-existing lesion model was compared with baseline and integrated models fitted on identical folds in the same sample: patients with WMH segmentations (n = 1,972) and patients with at least one pre-existing lesion segmentation (n = 1,973), respectively. For parsimony, only the baseline and integrated models of the WMH sample are shown; in the second sample, their mean AUPRC differed by at most 0.002. Violins show the distribution across the 100 outer folds, grey points individual folds, black diamonds the mean, and numbers above each violin the mean AUPRC.

AUPRC = area under the precision–recall curve; PSCI = post-stroke cognitive impairment; WMH = white matter hyperintensities.

**Text S1. Derivation of functional and structural lesion network mapping scores**

*Functional lesion network mapping.* Functional LNM scores estimate the brain-wide functional connectivity profile of the tissue affected by the infarct, using resting-state functional connectivity from healthy individuals as a normative reference.[1] Each patient's binary lesion mask, in MNI152 standard space, was used as a seed region in the Brain Genomics Superstruct Project 1000 (GSP1000) normative connectome, comprising resting-state fMRI data from 1,000 healthy participants. For each participant of GSP1000, the mean blood-oxygen-level-dependent timecourse across all voxels within the lesion mask was correlated with the timecourse of every other brain voxel, and the resulting correlation coefficients were Fisher r-to-z transformed. Voxelwise z-values were then combined across the 1,000 participants into a single lesion-seeded functional connectivity map by means of a one-sample t-test, yielding a t-statistic at each voxel that expresses how consistently that voxel is functionally connected to the lesioned tissue across the normative sample. The connectivity map was not thresholded. Functional LNM scores were obtained by averaging the t-statistics across all voxels of each of the 190 regions. Scores are therefore signed: positive values indicate that a region is positively coupled to the lesioned tissue in the normative connectome, and negative values indicate anticorrelation.

*Structural lesion network mapping.* Structural LNM scores estimate which white matter connections traverse the infarcted tissue, using tractography based on diffusion-weighted imaging of healthy individuals as a normative reference. Each patient's binary lesion mask was used to filter the dTOR985 normative tractogram, comprising whole-brain streamlines reconstructed from diffusion-weighted imaging in 985 healthy participants. We first identified the set of streamlines that intersected the lesion mask at any point along their trajectory. For each of the 190 regions, we then identified the subset of these lesion-intersecting streamlines associated with that region. The structural LNM score was defined as the proportion of the region's streamlines that intersected the lesion, expressed as a percentage. Scores are therefore bounded between 0 and 100 and normalised with respect to each region's own connectivity, such that a value of 0 indicates that no streamline associated with the region passes through the lesion and a value of 100 indicates that all of them do. Streamline–region association was determined by streamline endpoints for the cortical, subcortical, and cerebellar parcellations, and by streamline traversal for the XTRACT white matter tracts, reflecting the distinction between grey matter regions, which connections terminate in, and white matter tracts, which connections run through.

### Text S2. Algorithm comparison: XGBoost and TabPFN

To assess whether our findings depended on the choice of prediction algorithm, we re-estimated the integrated model with two alternative machine learning approaches: XGBoost,[2] a gradient-boosted decision-tree ensemble, and TabPFN,[3] a transformer-based tabular foundation model. Apart from the applied algorithm, the evaluation framework was identical to the main analysis.

Architecture. The stacked-ensemble architecture of the integrated model was retained unchanged; only the base learners of the three imaging views (lesion location, functional LNM, and structural LNM, each incorporating the baseline variables) were replaced by the respective backend. The demographics-only baseline learner remained the regularised linear (ridge) model in all configurations, so that the backend comparison specifically probes the modelling of the high-dimensional imaging representations. Feature preprocessing was identical across backends (median imputation and z-standardisation, fit within each training fold). As in the main analysis, base-learner predictions, i.e., predicted class probabilities for XGBoost and TabPFN and decision values for ridge, were generated out-of-fold by 10-fold inner cross-validation within each outer training fold, column-standardised, and combined by the same L2-regularised logistic regression meta-learner, whose regularisation constant was selected by 10-fold cross-validation on the out-of-fold prediction matrix.

XGBoost. Gradient-boosted trees were fit with a binary logistic objective, histogram-based tree construction, and the area under the precision–recall curve as the internal evaluation metric. Hyperparameters were fixed a priori rather than tuned per dataset: maximum tree depth 4, learning rate 0.05, subsample fraction 0.8, per-tree column sampling 0.8, minimum child weight 5, L2 regularisation $\lambda = 1.0$, L1 regularisation $\alpha = 0.5$, and a maximum of 500 boosting rounds. The only data-adaptive element was early stopping: within each fit, a stratified 15 % validation split was separated from the training data, and boosting was terminated after 50 rounds without improvement on this internal split. At no point did held-out test data enter model fitting or early stopping.

TabPFN. TabPFN was applied in its intended zero-shot, in-context mode: no gradient-based training or per-dataset hyperparameter tuning was performed. For each fit, the training data served as the context set and class probabilities for the evaluation data were obtained in a single forward pass with default inference settings on GPU hardware. Inference beyond the model's nominal pretraining regime was explicitly enabled where sample size or feature dimensionality required it.

Evaluation. All three backends were evaluated on identical outer folds under pooled 10 × 10 repeated stratified cross-validation, for all cognitive endpoints. Predictive performance was quantified as AUPRC.
